# Curvature-sensing peptide functions as a membrane interfactant that glues small extracellular vesicles to cell membranes and enhances vesicle cellular uptake

**DOI:** 10.1101/2025.08.06.669005

**Authors:** Kenichi Kawano, Kenta Hosokawa, Aoi Taniguchi, Yuuto Oosugi, Yuki Kuzuama, Katsumi Matsuzaki

## Abstract

Small extracellular vesicles (sEVs) are lipid nanoparticles secreted from mammalian cells that are involved in the transfer of informational or therapeutically effective substances between cells. Although the scope of research on sEVs as biocompatible carriers for drug delivery to diseased tissues and cells is expanding, several challenges remain, such as low cellular uptake efficiency and difficulty in loading drugs. It is desirable to alleviate the energy barriers associated with the cellular uptake of sEVs and minimize perturbation of sEVs when loading drugs onto them. In this study, we developed a simple drug-loading system for sEVs using a dimeric curvature-sensing peptide, which enhances sEV accumulation on the cell surface by acting as an adhesive, subsequently inducing endocytic uptake of sEVs through a clathrin-mediated pathway. The dimeric curvature-sensing peptide selectively binds to the sEV surface within 10 min, even in the presence of serum proteins, and functions as a membrane interfactant to reduce the energy barriers for the cellular uptake of sEVs. The cellular uptake of sEVs and the dimeric curvature-sensing peptide under coexisting conditions increased to over fivefold and 20-fold, respectively, compared with those administered alone. Furthermore, the dimeric curvature-sensing peptide can efficiently load anticancer drugs onto the surface of sEVs, and the system effectively induces apoptosis in two types of cancer cells. Dimeric curvature-sensing peptide is a novel technique with potential applications in drug delivery.

## Introduction

Small extracellular vesicles (sEVs) are biogenic lipid nanoparticles with diameters of 50 to 200 nm that are secreted from mammalian cells and play key roles in cell–cell communication between EV-donor and recipient cells^1^. Because they carry important biomarkers and signaling molecules involved in various human diseases, they hold great promise as pharmaceutical targeting vesicles. For instance, sEVs derived from mesenchymal stem cells (MSCs) have been reported to exhibit anticancer activities^2, 3^. The scope of research on sEVs is expanding to include the use of biomolecules that are effective in drug delivery and disease treatment. Chemical reactions^4, 5^, antibody recognition^6^, genetic engineering^6^, hydrophobic chain insertion^7, 8^, and electroporation^8, 9^ are widely used to load cargo into sEVs (Figure 1A). However, several challenges remain in the development of drug delivery systems, including difficulty in drug loading and low cellular uptake efficiency. The amount of drugs loaded onto sEVs using chemical reactions and antibody recognition depends on the expression levels of the target molecules. Tetraspanins, such as CD9, CD63, and CD81, are the most highly enriched human exosome marker proteins, and their mean expression levels are reported to be 12 molecules per vesicle (maximum: 138 molecules per vesicle)^10^. Genetic engineering is a powerful method for introducing functional tags into sEVs; however, the expression efficiency of tagged molecules in sEVs can be a limiting factor. The average numbers of CD9, CD63, and CD81 tagged with Halo-Tag7 (approximately 33 kDa) are estimated to be approximately 4–5 molecules per sEV^11^. Hydrophobic chain insertion and electroporation exhibit limitations in target selectivity and loading efficiency, respectively. Cellular uptake efficiency depends on particle concentration; however, sEVs inherently have a low capacity to induce uptake by recipient cells. Although the blood plasma concentration of EVs in healthy humans is estimated to be approximately 10^10^ particles/mL^12^, preparing highly concentrated sEVs loaded with therapeutic agents is labor-intensive and costly, as the yield from 10 mL of cultured mammalian cell media typically ranges from approximately 10^7^ to 10^9^ particles/mL, depending on the isolation method^13^. Moreover, because drug-loaded sEVs disperse in the bloodstream after injection, high particle concentrations are required in formulations to achieve therapeutic effects at target tissues. Thus, an alternative system capable of providing therapeutic efficacy even at lower particle concentrations is needed, particularly for *in vivo* applications. To address this challenge, we employed curvature-sensing peptides to enhance the therapeutic potential of sEVs.

**Figure 1.**
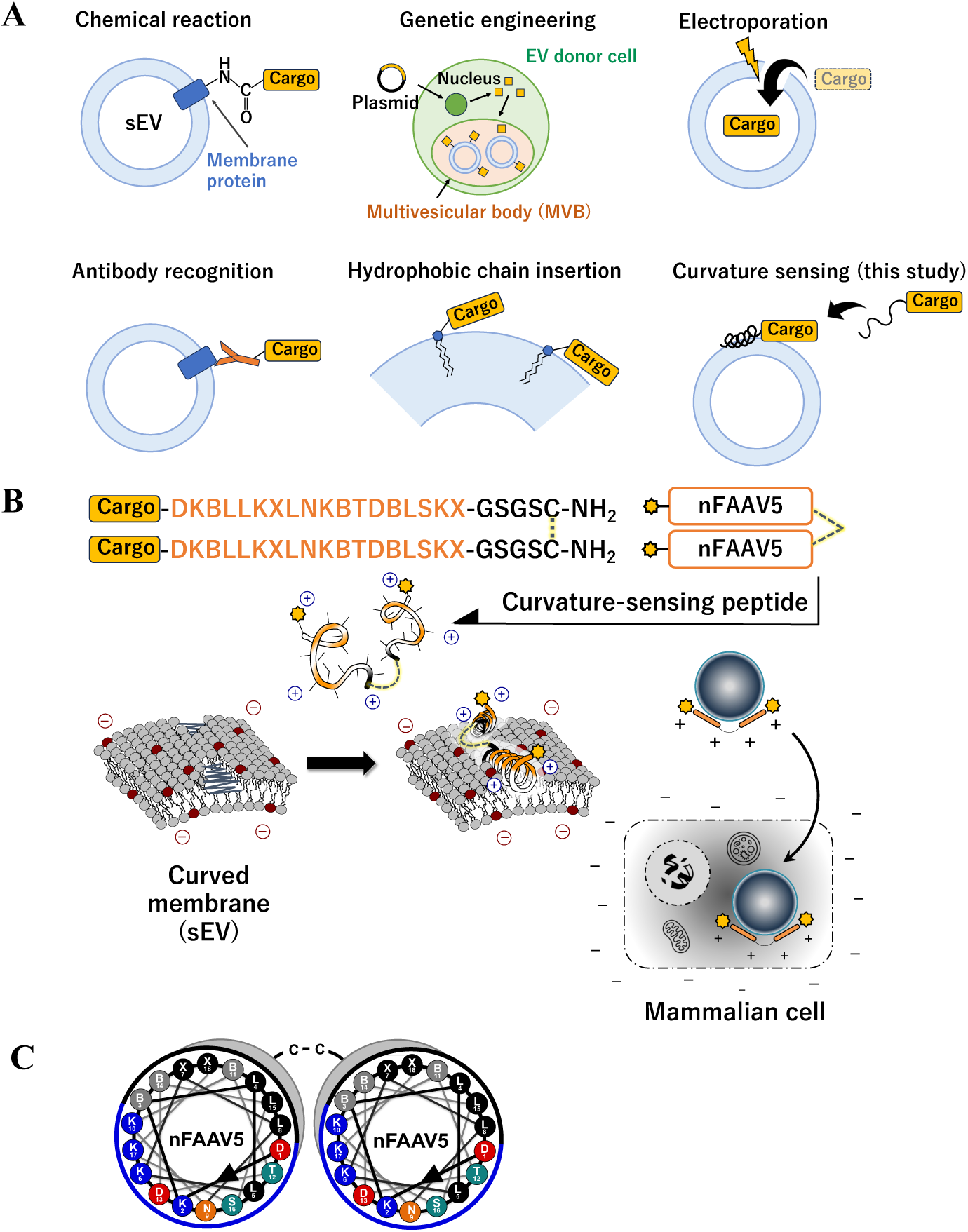
Conception of sEV modification. (A) Classification of methodologies for loading cargos on/in sEVs in conventional studies and the present study. (B) A schematic diagram of an sEV-modifying peptide binding to the sEV surface and enhancing cellular uptake of sEVs. (C) A helical wheel projection of the dimeric curvature-sensing peptide, di-nFAAV5 (sEV-modifying peptide).

Previously, we developed a biomarker-free EV-detecting curvature-sensing peptide, FAAV (named after four mutations: F8L, A11B, A11B, and V15L; B = 2-aminoisobutyric acid), based on the sequence of sorting nexin protein 1 of the Bin/amphiphysin/Rvs protein family^14^. The second-generation peptide, modified at the N-terminus and termed nFAAV5, has the ability to detect bacterial EVs within 5 min in cultured media, even in the presence of EV-secreting cells^15^. This capability enabled its application in simple, high-throughput screening of bacterial EVs from over 10,000 variant strains, leading to the identification of genes involved in bacterial EV production^16^. Furthermore, we developed a novel isolation system for mammalian sEVs—EV-CaRiS (EV catch-and-release isolation system)^13^—using a net charge–invertible curvature-sensing peptide (NIC4) based on nFAAV5.

In the present study, we propose a novel strategy for enhancing the cellular uptake of sEVs and a simple drug-loading system for sEVs using a curvature-sensing peptide (Figure 1A), which recognizes lipid-packing defects by inserting its hydrophobic face of the amphipathic α-helix into the curved membrane^17, 18^ (Figures 1B and C). Curvature-sensing peptides coat the membrane surface of sEVs, similar to a net wrapped around a football. We utilized this property of curvature-sensing peptides to develop a concept that enables efficient loading of therapeutic agents onto the sEV surface. Herein, we designed a novel dimeric nFAAV5 (di-nFAAV5) as an sEV-modifying peptide with higher affinity for sEVs than conventional peptides. It exhibited a remarkable enhancement in the cellular uptake of sEVs, even in the presence of serum proteins, by over fivefold through the clathrin-mediated endocytosis pathway, subsequently enabling the effective delivery of apoptosis-inducing peptide drugs to cancer cells by over 20-fold. We found that di-nFAAV5 functions as a *membrane interfactant* to enhance the attachment of sEVs to the plasma membrane of cells, acting as an adhesive, and reduces the energy barrier for the cellular uptake of sEVs. Here, “*membrane interfactant*” is a portmanteau of “membrane interface” and “surfactant.” It activates interactions between membrane interfaces but does not disrupt or solubilize membranes, nor does it exhibit cytotoxicity. This property of curvature-sensing peptides may contribute to the development of novel drug delivery systems using sEVs as carriers.

## Materials and Methods

### Peptide synthesis

Peptides were synthesized according to previously described procedures^13^. Based on Fmoc solid-phase peptide chemistry, Fmoc-amino acids (3 equiv.), coupling reagent (3 equiv.), and base (6 equiv.) were used to elongate the main chain on an H-Rink ChemMatrix amide resin (1 equiv.) (Sigma-Aldrich, St. Louis, MO). The coupling reaction was performed at 25 °C for 1 h using 1-[bis(dimethylamino)methylene]-1*H*-benzotriazolium 3-oxide hexafluorophosphate, 1-hydroxybenzotriazole, and *N,N*-diisopropylethylamine in *N,N*-dimethylformamide (DMF). Fmoc-Lys(Mtt)-OH was used to label nitrobenzoxadiazole (NBD) dye at the N- or C-termini of **3**–**7**. Fmoc-Cys(Trt)-OH coupling was performed using Oxyma Pure/diisopropylcarbodiimide in DMF. Fmoc-protected *S*-pentenylalanine (Z), *S*-octenylalanine (Z′), and 2-aminoisobutyric acid (B), along with subsequent amino acids, were coupled at 40 °C for 2 h using (1-cyano-2-ethoxy-2-oxoethylidenaminooxy)dimethylamino-morpholinocarbenium hexafluorophosphate and *N,N-* diisopropylethylamine in DMF. For NBD labeling, 4-fluoro-7-nitrobenzofurazan (2.0 equiv.) (Dojindo, Kumamoto, Japan) was reacted on resin with the N-terminal main chain or N- or C-terminal Lys side chains in DMF for 16 h at 25 °C after removal of the Mtt group in 20% hexafluoroisopropanol^14^. Deprotection and cleavage of peptides from the resin were conducted in a cocktail of trifluoroacetic acid/water/triisopropylsilane/1,2-ethanedithiol (94:2.5:1.0:2.5, v/v) for 3 h at 25 °C. The peptides were purified by RP-HPLC on a 5C_18_ column (4.6 mm I.D. × 150 mm) from COSMOSIL (Nacalai Tesque, Kyoto, Japan) or InertSustainSwift (GL Sciences, Tokyo, Japan) using a linear gradient from 30% to 80% acetonitrile in 0.1% aqueous trifluoroacetic acid for 20 min at 40 °C at a flow rate of 1.0 mL/min. Mass was confirmed by electrospray ionization mass spectrometry using an LCMS-8040 mass spectrometer (Shimadzu, Kyoto, Japan). Dimerization of the peptides was performed in 10 mM Tris buffer (pH 6.0) containing 50% (v/v) dimethyl sulfoxide^19^ for 16–18 h at 25 °C. Ring-closing metathesis was performed using AquaMet catalyst (0.3 equiv.) (Tokyo Chemical Industry, Tokyo, Japan) and MgCl₂·6H₂O (400 equiv.) in a solution of 160 mM phosphate buffer containing 4.8 M guanidine hydrochloride and DMF (4:1, v/v) for 2 h at 60 °C^20^. The peptides were purified by RP-HPLC. Absorbance spectra were measured in pH-controlled phosphate-buffered saline [PBS(–): 137 mM NaCl, 2.7 mM KCl, 8.1 mM Na₂HPO₄, and 1.5 mM KH₂PO₄; pH 7.4] using an ultraviolet-visible spectrophotometer (UV-2550; Shimadzu). The concentrations of NBD-labeled peptides were calculated based on the absorption of NBD at 467 nm (ε = 28,000 cm⁻¹ M⁻¹). Because dimerized peptides **2**–**7** contained two NBD dyes per molecule, their net concentrations were determined by dividing the NBD absorbance by two. The concentrations of nonlabeled peptides were estimated based on their molecular weights.

### Liposome preparation

Liposomes were prepared as previously described^13^. Non-labeled liposomes composed of 1-palmitoyl-2-oleoyl-*sn*-glycero-3-phosphocholine (POPC), cholesterol, and 1-palmitoyl-2-oleoyl-*sn*-glycero-3-phospho-L-serine (POPS) (75:15:10 mol%) were used in the peptide-binding assay. ATTO647N-labeled liposomes composed of POPC, cholesterol, POPS, and 1,2-dipalmitoyl-*sn*-glycero-3-phosphoethanolamine labeled with Atto 647N (ATTO647N-DPPE) (75:15:10:0.1 mol%) were used for the cellular uptake assay. Lipid concentrations were determined by phosphorus analysis^21^.

### Measurement of particle sizes and ζ-potential values

Particle sizes were measured using a dynamic light scattering system (Zetasizer Nano ZS, Malvern Panalytical, Worcestershire, UK) and a nanoparticle tracking analysis system (NanoSight 3000, Malvern Panalytical) at 25 °C. ζ-Potential values were determined using the Zetasizer Nano ZS at 25 °C with disposable cells (DTS1070).

### Determination of *K_d_* values

NBD-labeled peptides at a final concentration of 0.5 µM were mixed with liposomes (diameter: 90.5 ± 1.1 nm; ζ-potential: –28.0 ± 0.6 mV) at lipid concentrations of 0, 2, 5, 10, 20, 50, 100, and 200 µM in 150 µL/well PBS(–) in a 96-well plate (As One, Osaka, Japan). NBD fluorescence intensity was measured after incubation for 10 min at 25 °C using a plate reader, EnVision (PerkinElmer Japan, Yokohama, Japan), at Ex. 480/30 nm and Em. 535/25 nm. After subtraction of the peptide background, *K_d_* values were determined using Eq. (1), based on the Langmuir isotherm model^18^:

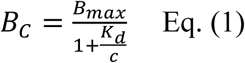

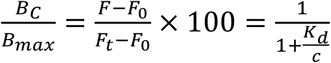

where *B_c_*, *B_max_*, and *c* represent the vesicle-binding level at each point, maximum binding level, and the given lipid concentration of vesicles, respectively. The ratio of *(F – F_0_)* / *(F_t_ – F_0_)* was calculated and plotted as a function of *c*, where *F*_0_ and *F* denote the fluorescence intensities of NBD-peptides in the absence and presence of vesicles, respectively. The free peptide fraction at each lipid concentration was estimated using the following equation: (1 – *B_c_* / *B_max_*).

### Circular dichroism (CD) spectroscopy

Peptides (40 µM) and liposomes (2 mM) were mixed in 400 µL of Dulbecco’s PBS^15^ and incubated for 10 min at 25 °C. CD spectra were obtained using a quartz cuvette with a light path length of 1 mm and J-820 CD spectrometer (JASCO, Tokyo, Japan). The final spectrum was the average of five scans after subtraction of the blank spectra of the buffer or liposomes under identical conditions.

### Cell culture

HeLa and PANC-1 cells were cultured in Dulbecco’s modified Eagle’s medium (DMEM) (high glucose, 4.5 g/L) containing 10% (v/v) heat-inactivated fetal bovine serum (FBS) at 37 °C with 5% CO_2_ in a humidified incubator. MSC-R37 cells were obtained from RIKEN BioResource Research Center (Cell #: HMS0020) (RIKEN BRC, Ibaraki, Japan)^22^ and cultured in DMEM (low glucose, 1.0 g/L) supplemented with 3 ng/mL basic fibroblast growth factor (bFGF) and 10% (v/v) heat-inactivated FBS at 37 °C with 5% CO_2_.

### sEV isolation and 1,1′-dioctadecyl-3,3,3′,3′-tetramethylindodicarbocyanine (DiD) labeling

At 24 h after seeding MSC-R37 cells in 10 mL DMEM (low glucose) containing 3 ng/mL bFGF and 10% (v/v) FBS in a 10 cm culture dish, the cells were washed five times with 2 mL of fresh medium. The cells were then incubated for an additional 24 h in 10 mL DMEM (low glucose) supplemented with 3 ng/mL bFGF and 10% (v/v) exosome-free FBS. The cell-cultured medium (10 mL) was collected, and sEVs were isolated according to our previous study^13^. Labeling of sEVs isolated from MSC-R37 cells was performed using DiD at a concentration of 2 µM at 37 °C for 20 min (DiD-MSC-sEVs). The free dye was removed as previously described^23^.

### Western blotting

Western blotting was performed according to our previous study^13^.

### Observation of cellular uptake by confocal laser scanning microscopy (CLSM)

HeLa and PANC-1 cells were seeded at a density of 2.5 × 10^4^ cells/compartment (total: 1.0 × 10^5^ cells/dish) in a 35 mm glass-bottom dish with four compartments (CELLview, Advanced TC-treated) (Greiner Bio-One, Frickenhausen, Germany). After culture at 37 °C for 24 h, the cells were treated with NBD-labeled peptides (1–2 µM) and either ATTO647N-liposomes (20 µM lipid concentration) or DiD-MSC-sEVs (2.0 × 10^8^ particles/mL) in DMEM (high glucose, 4.5 g/L) containing 10% (v/v) exosome-free FBS [DMEM(+)] (250 µL/compartment). Time-course dual-color imaging of live cells was performed using Ex. 488 nm/Em. 500–550 nm and Ex. 638 nm/Em. 663–738 nm by CLSM on an A1Rsi microscope (Nikon, Tokyo, Japan) equipped with a heated stage at 37 °C and 5% CO_2_ supply. Images were acquired at 3 min intervals for 1 h immediately following treatment with **2** and DiD-MSC-sEVs. Internalization was observed after incubation for more than 1 h at 37 °C in a 5% CO_2_ incubator. Colocalization data were analyzed using Fiji software (ImageJ 1.54f).

### Colocalization analysis of dual-color confocal images

Colocalization of **2** and DiD-MSC-sEVs was analyzed according to Eq. (2), as described in a previous study^24^:

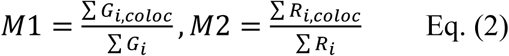

where *G*_i, coloc_ and *G*_i_ denote green signals (NBD) colocalized with red signals (DiD) and arbitrary green signals in the region of interest in the green channel, respectively. *R*_i, coloc_ and *R*_i_ denote red signals (DiD) colocalized with green signals (NBD) and arbitrary red signals in the region of interest in the red channel, respectively. The coefficients M1 and M2 range from zero to one, where *G*_i, coloc_ = *G*_i_ if *R*_i_ > 0, and *G*_i, coloc_ = 0 if *R*_i_ = 0 (M1); *R*_i, coloc_ = *R*_i_ if *G*_i_ > 0, and *R*_i, coloc_ = 0 if *G*_i_ = 0 (M2). These two coefficients are defined as parameters independent of signal intensity.

### Ca²⁺ response assay

HeLa and PANC-1 cells were seeded into 35 mm glass-based dishes (Iwaki, Shizuoka, Japan) at a density of 1 × 10^5^ cells/dish in DMEM (high glucose) containing 10% (v/v) FBS. For stimulation with 100 µM ATP, the Ca^2+^ response assay was performed according to our previous report^25^. For stimulation with **2** or MSC-sEVs, the cells were washed twice with DMEM(+) containing 0.36 mg/mL probenecid. Rhod-2 AM (Setareh Biotech, Tyler, TX) was then loaded into the cells at a final concentration of 5 µM in DMEM(+) containing 0.36 mg/mL probenecid at 37 °C for 1 h. The cells were washed with DMEM(+) containing 0.36 mg/mL probenecid twice. The fluorescence intensity of Rhod-2 was recorded for 600 s at 20 s intervals immediately after the addition of **2** and MSC-sEVs at final concentrations of 1 µM and 2 × 10^8^ particles/mL, respectively, in DMEM(+) containing 0.36 mg/mL probenecid. Time-course observations were performed using CLSM at Ex. 488 nm/Em. 500–550 nm and Ex. 561 nm/Em. 570–620 nm on a heated stage at 37 °C with a 5% CO₂ supply.

### Inhibition assay of cellular uptake

To inhibit cell membrane accumulation, cells were treated in DMEM(+) containing **2** (2 µM), DiD-MSC-sEVs (2 × 10^8^ particles/mL), and heparin (100 or 500 µg/mL)^8, 26^ for more than 1 h at 37 °C in a 5% CO₂ incubator. To inhibit membrane curvature induction, cells were treated in HEPES buffer [20 mM HEPES, 25 mM glucose, 1 mM MgCl₂, 2 mM CaCl₂ (pH 7.4)] containing either 150 mM NaCl (isotonic buffer) or 70 mM NaCl and 30 mM KCl (hypotonic buffer), in the presence of **2** (2 µM) and DiD-MSC-sEVs (2.0 × 10^8^ particles/mL) for more than 1 h at 37 °C in a 5% CO_2_ incubator, according to the reported study^27^. To inhibit internalization, the cells were preincubated in DMEM(+) containing inhibitors (EIPA: 10 µM; wortmannin: 1 µM; cytochalasin D: 200 nM; pitstop2: 15 µM; dynasore: 15 µM)^28, 29^ at final concentrations for 30 min at 37 °C in a 5% CO_2_ incubator. The medium was then replaced with DMEM(+) containing **2** (1 µM), DiD-MSC-sEVs (1.0 × 10^8^ particles/mL), and the same inhibitors at identical concentrations. In the absence of **2**, DiD-MSC-sEVs were used at 1.0 × 10^9^ particles/mL. After incubation for 1 h at 37 °C in a 5% CO_2_ incubator, the medium was removed, and fresh DMEM(+) was added for uptake observation. Pitstop2, a poorly water-soluble compound, was preincubated for more than 15 min at 60 °C in medium to improve solubility, followed by cooling to 25°C before use. Dynasore can be inactivated by serum proteins^30^; therefore, FBS pretreated by ultrafiltration using an Amicon Ultra-15 centrifugal filter with a 30k molecular weight cutoff (Merck) was used.

### Anticancer activity and caspase-8 assays

Anticancer activity was determined using the WST-8 reagent with the Cell Counting Kit-8 (Dojindo, Kumamoto, Japan). HeLa and PANC-1 cells were seeded at a density of 5 × 10^3^ cells/well in a 96-well microplate (Iwaki) in DMEM containing 10% (v/v) FBS and cultured for 24 h at 37 °C. For the cytotoxicity assay, cells were treated with DMEM(+) or PBS(+) [137 mM NaCl, 8.1 mM Na₂HPO_4_, 2.68 mM KCl, 1.47 mM KH_2_PO_4_, 0.33 mM MgCl_2_, and 0.90 mM CaCl_2_, pH 7.4) in the presence of **2′** (1 µM) and liposomes (30 µM). For the anticancer activity assay, MSC-sEVs (6.3 × 10⁹ particles/mL) were preincubated with peptide **8** (1 µM) in DMEM(+) for 10 min at 25 °C (**solution A**); streptavidin (SA) (2 µM) was preincubated with anticancer peptides **9** or **10** (6 µM) in DMEM(+) for 10 min at 25 °C (**solution B**). Solutions **A** and **B** were then combined to form the sEV complex as illustrated in Figure 5D. Cells were treated with the complex after being washed with 100 µL/well PBS(–). After 1 h of incubation at 37 °C in a 5% CO₂ incubator, the medium was replaced with fresh high-glucose DMEM(+) followed by a further 23 h incubation. After removing the medium, cells were treated with 100 µL/well of high-glucose DMEM containing 10% (v/v) FBS and 5% (v/v) WST-8 and incubated for 1 h (HeLa) or 2.5 h (PANC-1) at 37 °C in a 5% CO₂ incubator. The absorbance of the WST-8 formazan was measured at 450 nm using an EnVision2105 multimode plate reader (PerkinElmer). Cell viability was calculated based on the absorbance of WST-8 formazan in untreated healthy cells, after subtraction of background signals from the medium. Caspase-8 activity assay was performed using the Cell Meter™ Caspase-8 Activity Apoptosis Assay Kit (AAT Bioquest, Pleasanton, CA) with minor modifications. The cells were treated with the sEV complex solution in the same manner and incubated at 37 °C for 23 h in fresh DMEM(+). A mixture of anti-Fas antibody (500 ng/mL) and cycloheximide (500 ng/mL) was used as the positive control (incubation time: 24 h). The substrate (Ac-IEAD)-AMC, an indicator of caspase-8 activity, was added directly to the cultured medium (320-fold dilution). AMC fluorescence was measured at 450 nm using a Wallac 1420 Victor2 microplate reader (PerkinElmer) at Ex. 380 nm after incubation at 37 °C for 30 min. The background signal from the medium was subtracted from the AMC fluorescence of the cells. A cell membrane perturbation assay was performed using the membrane-impermeable dye propidium iodide (PI) at a final concentration of 3.5 µM. After incubation with PI at 37 °C for 30 min, CLSM images of the PI-stained cells were acquired at Ex. 561 nm. The cells treated with 100% MeOH were used as positive controls.

## Results

### Design and functions of sEV-modifying peptides

We found that a dimeric curvature-sensing peptide labeled with an NBD dye at the N-terminus (di-NBD-nFAAV5, entry **2**) (Table 1) exhibited broad applicability as an sEV-modifying peptide. The drug-loading efficiency of sEVs depends on the number of sEV-binding peptides. Monomeric nFAAV5 labeled with NBD (entry **1**) (Table 1) achieved selective binding to EVs within 5 min by simply placing it in cultured media without separation steps^15^; di-nFAAV5 was designed to improve affinity for sEVs and plateau binding levels because the plateau level in the binding curves is proportional to the number of peptides on the sEVs; di-nFAAV5 has two sEV-binding regions, allowing it to maintain a longer bound state than the monomeric one because one region dissociates from sEVs while the other remains bound. NBD, a hydrophobic environment-sensitive dye that emits strong fluorescence in membranes, was conjugated with peptides instead of a drug. To optimize the labeling position, two dimeric peptides labeled with NBD at the N-terminus (entry **2**) or C-terminus (entry **3**) were prepared (Table 1 and Figures S1–S3). NBD and DiD (for labeling sEVs) were selected as pairs that do not cause optical interference, such as fluorescence resonance energy transfer.

**Table 1.** sEV-modifying peptide candidates and stapled anticancer peptides.

| Entry | Peptide Name | Sequence<sup>(a)</sup> | Theoretical value[M+H]<sup>+</sup> | Calculated m/z |
| --- | --- | --- | --- | --- |
| 1 | NBD-nFAAV5 | NBD-DKBLLKXLNKBTDLSKX-GSGSK-NH<sub>2</sub> | 2575.46 | 2575.47 |
| 2 | di-NBD-nFAAV5 | NBD-DKBLLKXLNKBTDLSKX-GSGS-C-NH<sub>2</sub><br>NBD-DKBLLKXLNKBTDLSKX-GSGS-C-NH<sub>2</sub> | 5097.72 | 5098.74 |
| 3 | di-nFAAV5-NBD | DKBLLKXLNKBTDLSKX-GSGSK(NBD)-C-NH<sub>2</sub><br>DKBLLKXLNKBTDLSKX-GSGSK(NBD)-C-NH<sub>2</sub> | 5353.91 | 5354.94 |
| 4 | di-K(NBD)-nFAAV5 | K(NBD)-DKBLLKXLNKBTDLSKX-GSGS-CK-NH<sub>2</sub><br>K(NBD)-DKBLLKXLNKBTDLSKX-GSGS-CK-NH<sub>2</sub> | 5610.10 | 5612.11 |
| 5 | di-K(NBD)-nFAAV6 | K(NBD)-DRBLLRXLNRBTDLSRX-GSGS-CK-NH<sub>2</sub><br>K(NBD)-DRBLLRXLNRBTDLSRX-GSGS-CK-NH<sub>2</sub> | 5834.15 | 5836.06 |
| 6 | di-C8-K(NBD)-nFAAV5 | C8-K(NBD)-DKBLLKXLNKBTDLSKX-GSGS-CK-NH<sub>2</sub><br>C8-K(NBD)-DKBLLKXLNKBTDLSKX-GSGS-CK-NH<sub>2</sub> | 5862.31 | 5864.28 |
| 7 | di-C8-K(NBD)-nFAAV6 | C8-K(NBD)-DRBLLRXLNRBTDLSRX-GSGS-CK-NH<sub>2</sub><br>C8-K(NBD)-DRBLLRXLNRBTDLSRX-GSGS-CK-NH<sub>2</sub> | 6086.38 | 6088.39 |
| 8 | di-biotin-nFAAV5 | biotin-DKBLLKXLNKBTDLSKX-GSGS-C-NH<sub>2</sub><br>biotin-DKBLLKXLNKBTDLSKX-GSGS-C-NH<sub>2</sub> | 5223.87 | 5222.91 |
| 9 | biotin-SAH-p53-4 | biotin-LSQETFZ'DLWKLLZEN-NH<sub>2</sub> | 2253.20 | 2253.19 |
| 10 | biotin-BIM-BH3 | biotin-IWIAQELRRIGDEFNZYYAZR-NH<sub>2</sub> | 2888.50 | 2889.26 |

<sup>a</sup>) The single letters of Z, Z', X, and B indicate (S)-2-(4-pentenyl)alanine, (R)-2-(7-octenyl)alanine, norleucine, and 2-aminoisobutyric acid, respectively. Norleucine was used instead of methionine to prevent the oxidation. C8, C-C, and Z-Z' are an octanoic acid, disulfide bond, and the crosslinking of stapled peptides, respectively. The C-termini of all peptides were amidated.

Predictably, **2** exhibited the highest affinity (dissociation constant, *K*_d_ = 3.5 µM at a lipid concentration; [lipids (3 µM)] ≈ [vesicles (3.6 × 10⁹ particles/mL)]^13^) for sEV-mimicking liposomes (Figure 2A), and this affinity was 15.7-fold higher than that of **1**. Conversely, **3** showed a remarkably reduced affinity (*K*_d_ = 258.9 µM), indicating that dye labeling at the C-terminal hinge region may impair vesicle binding of the dimeric peptide. The amount of **2** bound to sEVs was estimated to be 3,068 peptides/vesicle at the plateau level, based on previous data for **1** (1,361 peptides/vesicle^31^). Note that **2** (dimeric form) was used at half the concentration of **1** (monomeric form); thus, the ratio of fluorescence intensities was proportional to the number of peptides bound to vesicles. At the sEV secretion level in cultured media (10^8^–10^9^ particles/mL)^13^, the number of **2** peptides bound per vesicle was estimated to be approximately 140–590 based on Eq. (1), which is significantly higher than the reported expression levels of exosome marker proteins (4–12 molecules/vesicle)^10, 11^. The CD spectrum measurement revealed that **1** changes its secondary structure from a random coil to an amphipathic α-helix (Figure 2B). By contrast, **2** does not appear to change its structure (Figure 2B), although an increase in NBD fluorescence intensity was observed in the presence of vesicles (Figure 2A). These data suggest that **2** undergoes a transition from a closed, leucine zipper-like form (Figure 1C) in solution to an open form suited for vesicle binding. To observe the cellular uptake of sEVs in the presence of **2**, MSC-sEVs isolated by EV-CaRiS^13^ were labeled with the membrane stain DiD, with unbound dye removed according to a previously reported method^23^. Similar particle size distributions were observed between non-labeled and DiD-labeled MSC-sEVs (Figure 2C). Immunoblotting and fluorescence bands for the exosome biomarker CD63 and DiD, respectively, were detected simultaneously for DiD-MSC-sEVs (Figure 2C), indicating successful labeling. The ζ-potentials of non-labeled and DiD-labeled MSC-sEVs were –13.5 and +7.2 mV, respectively; a slight increase in ζ-potential was observed in the presence of non-labeled di-nFAAV5 (**2′**) (Figure 2D and S2). Importantly, we found that the enhanced cellular uptake of sEVs described subsequently is achieved by the function of **2** but is independent of the ζ-potential or DiD labeling of sEVs (Figures 2E and F). Although no remarkable intracellular signal of DiD-MSC-sEVs was observed in the absence of **2** (Figure 5A), a high accumulation of DiD-MSC-sEVs on the surface was observed in the presence of **2** using time-lapse imaging (Figure 2E). High colocalization coefficients at each time point (Figure 2E) demonstrated that **2** and DiD-MSC-sEVs accumulated together on the cell surface, even in the presence of FBS. For non-labeled MSC-sEVs, we evaluated the enhanced cellular accumulation by monitoring Ca^2+^ responses^11^. ATP treatment was used as a positive control to stimulate the ATP receptors expressed on the cell surface^32, 33^. Although the Ca^2+^ response level was not significantly different between MSC-sEVs treatment with and without **2**, there was a large difference in the detected frequency of Ca^2+^ responses (Figure 2F; “*n*” indicates the number of spikes reflecting Ca^2+^ responses). This revealed that non-labeled MSC-sEVs in the presence of **2** enhanced Ca^2+^ responses in HeLa and PANC-1 cells. Notably, cell binding of **2** alone was not observed (Figures 2F and 5A), indicating that **2** did not accumulate on the cell surface in the absence of sEVs.

**Figure 2.**
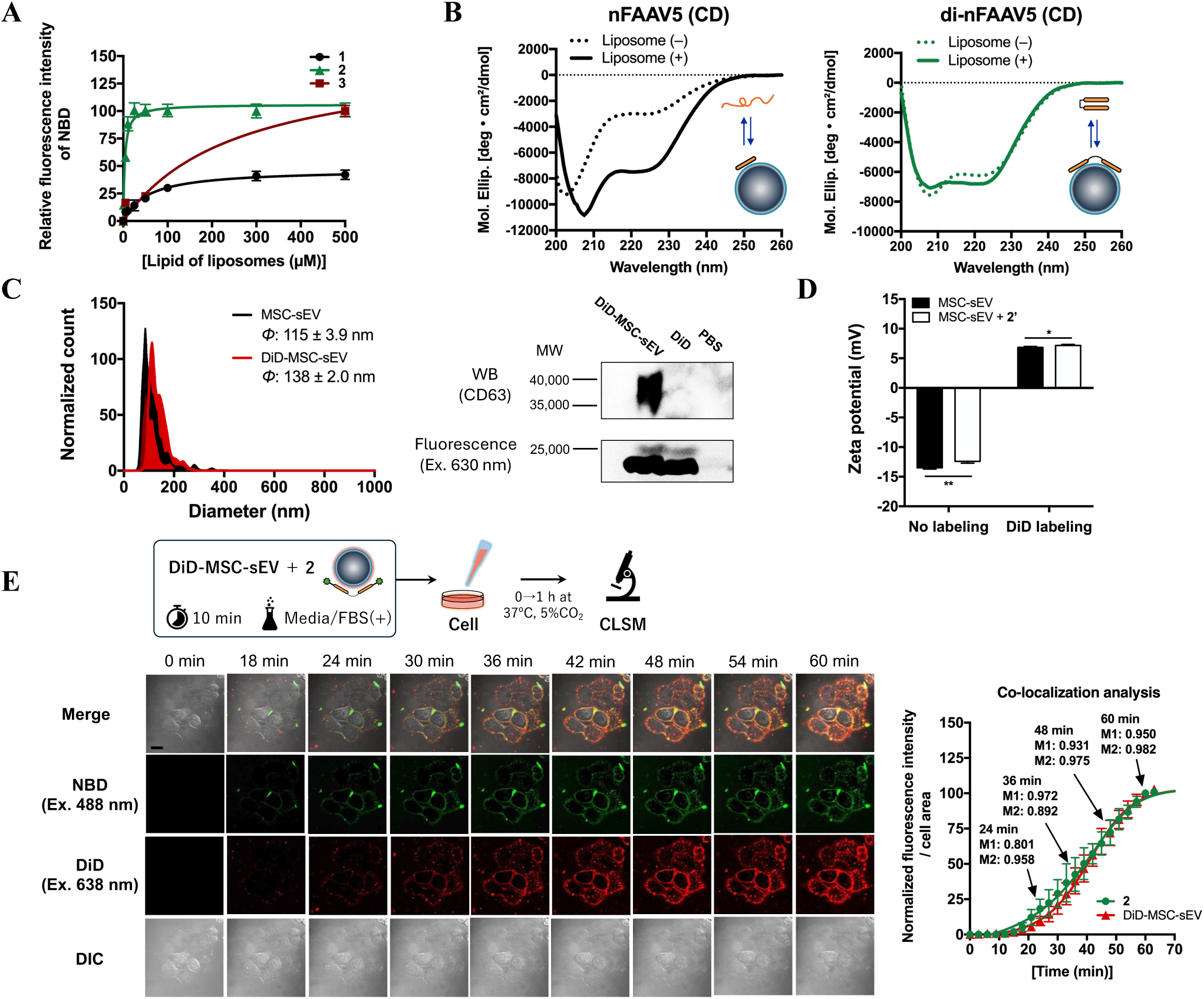
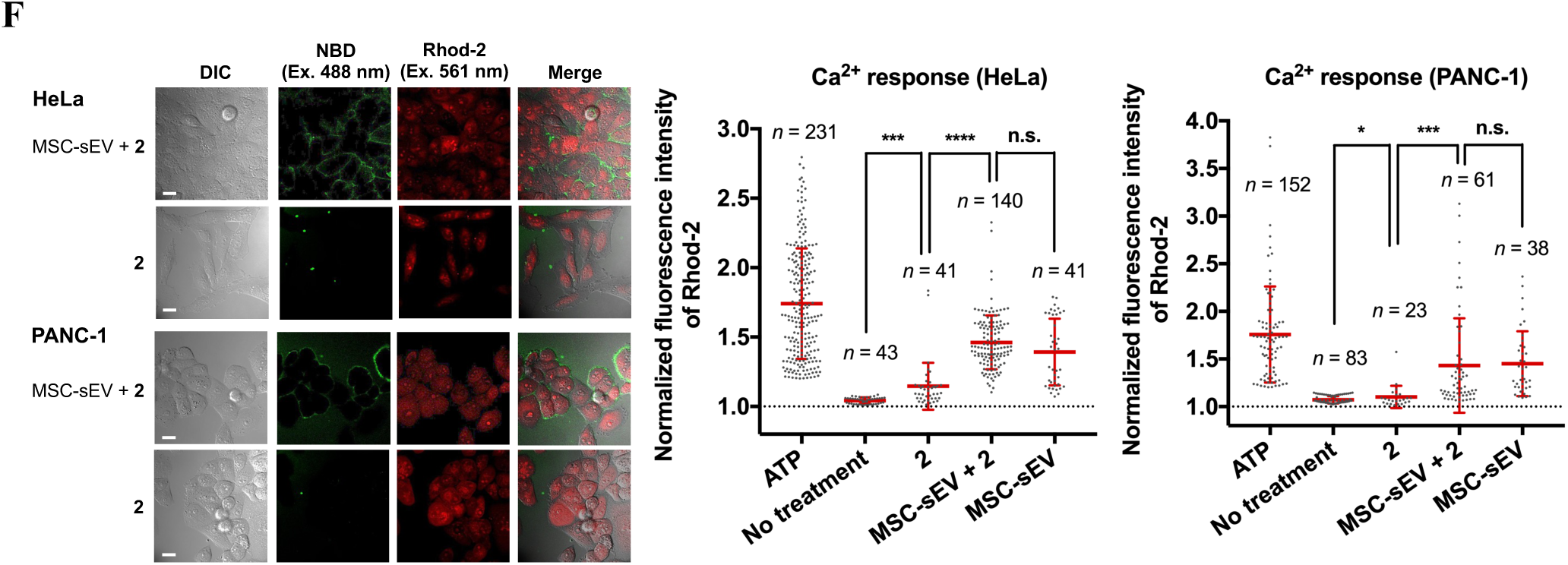
Basal properties of curvature-sensing peptides and isolated sEVs. (A) Relative fluorescence intensities of the NBD dye of **1**–**3** were plotted as a function of lipid concentration of liposomes with a diameter of approximately 91 nm (POPC:cholesterol:POPS = 75:15:10 mol%) (mean ± S.D., *n* = 3). The *K*_d_ values of **1** to **3** were calculated as 54.9, 3.5, and 258.9 µM in lipid concentration of liposomes based on Eq. (1), respectively. (B) CD spectra of non-labeled peptides in the absence and presence of liposomes. (C) Particle size distributions of non- and DiD-labeled sEVs isolated from MSC-R37 cells. The median diameters of non- and DiD-labeled sEVs were 115 ± 3.9 and 138 ± 2.0 nm, respectively (mean ± S.D., *n* = 5). Immunoblotting and fluorescent bands for CD63 and DiD (Ex. 630 nm) were imaged. Loading amount of sEVs: 2.0 × 10^9^ particles/lane. (D) Zeta potentials of non- and DiD-labeled sEVs in the absence and presence of **2** (mean ± S.D., *n* = 5). Statistical analysis was performed using Student’s *t*-test at an alpha level of 0.05 (\**p* < 0.05, \*\**p* < 0.01). (E) Time-lapse imaging of DiD-MSC-sEV (2 × 10^8^ particles/mL) accumulation on the surface of PANC-1 cells in the presence of **2** (2 µM) at 3 min intervals for 60 min at 37 °C in media/FBS(+). The scale bar is 10 µm. Time-courses of normalized fluorescence intensities of NBD and DiD per cell area are shown at right (mean ± S.D., *n* = 10 cells). Colocalization analysis of dual-color confocal images was performed using Eq. (2); the colocalization coefficients M1 (proportion of NBD signals colocalized with DiD signals) and M2 (proportion of DiD signals colocalized with NBD signals) at 24, 36, 48, and 60 min are shown. The closer the coefficient is to one, the higher the degree of colocalization. (F) Ca^2+^ responses of HeLa and PANC-1 cells treated with **2** in the presence or absence of non-labeled MSC-sEVs using the Ca^2+^ indicator Rhod-2. The “*n*” value indicates the number of spikes observed in Rhod-2 fluorescence time-courses (10 cells used for analysis).

### Elucidation of mechanism underlying sEV accumulation on the cell surface and uptake in the presence of an sEV-modifying peptide

We found that **2** bound to sEVs enhanced their accumulation on the cell surface as an adhesive through electrostatic interactions with negatively charged molecules, such as heparan sulfate proteoglycans^34^ and phosphatidylserine (PS) head groups of lipids, followed by membrane curvature induction prior to internalization (Figure 3A). To elucidate the mechanism of enhanced accumulation of sEVs on the cell surface, membrane binding of DiD-MSC-sEVs and **2** was observed in the presence of heparin. Heparin is widely used to prevent the attachment of positively charged cell-penetrating peptides to the cell surface^35, 36^. Heparin treatment significantly reduced the fluorescent signals of both DiD and NBD on the cell surface but did not influence the fluorescence intensity of NBD in the presence of MSC-sEVs in solution (Figure 3B), demonstrating that **2** maintains an sEV-bound state through hydrophobic interactions; however, its electrostatic interaction with the cell surface was abolished by heparin treatment (Figure 3A). Subsequently, to elucidate the mechanism underlying the enhanced uptake of sEVs, the internalization of DiD-MSC-sEVs and **2** was observed in isotonic or hypotonic buffers. Under hypotonic conditions, increased membrane stretching prevents internalization^27^. As expected, DiD-MSC-sEVs coated with **2** attached to the cell surface, but internalization was completely prevented under hypotonic conditions; conversely, both attachment and internalization were observed under isotonic conditions (Figure 3C). These results demonstrate that deformation of the plasma membrane is involved in the cellular uptake of sEVs in the presence of **2**, and **2** functions as an adhesive to enhance sEV accumulation on the cell surface and enables effective cellular uptake.

**Figure 3.**
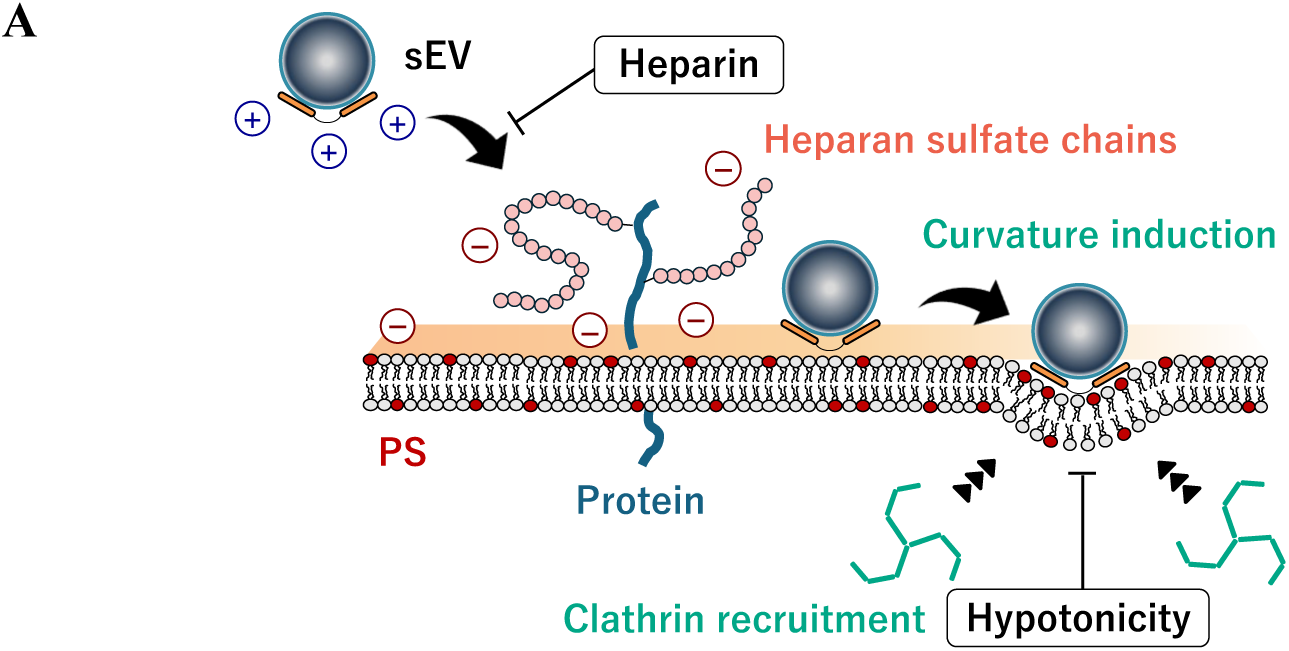
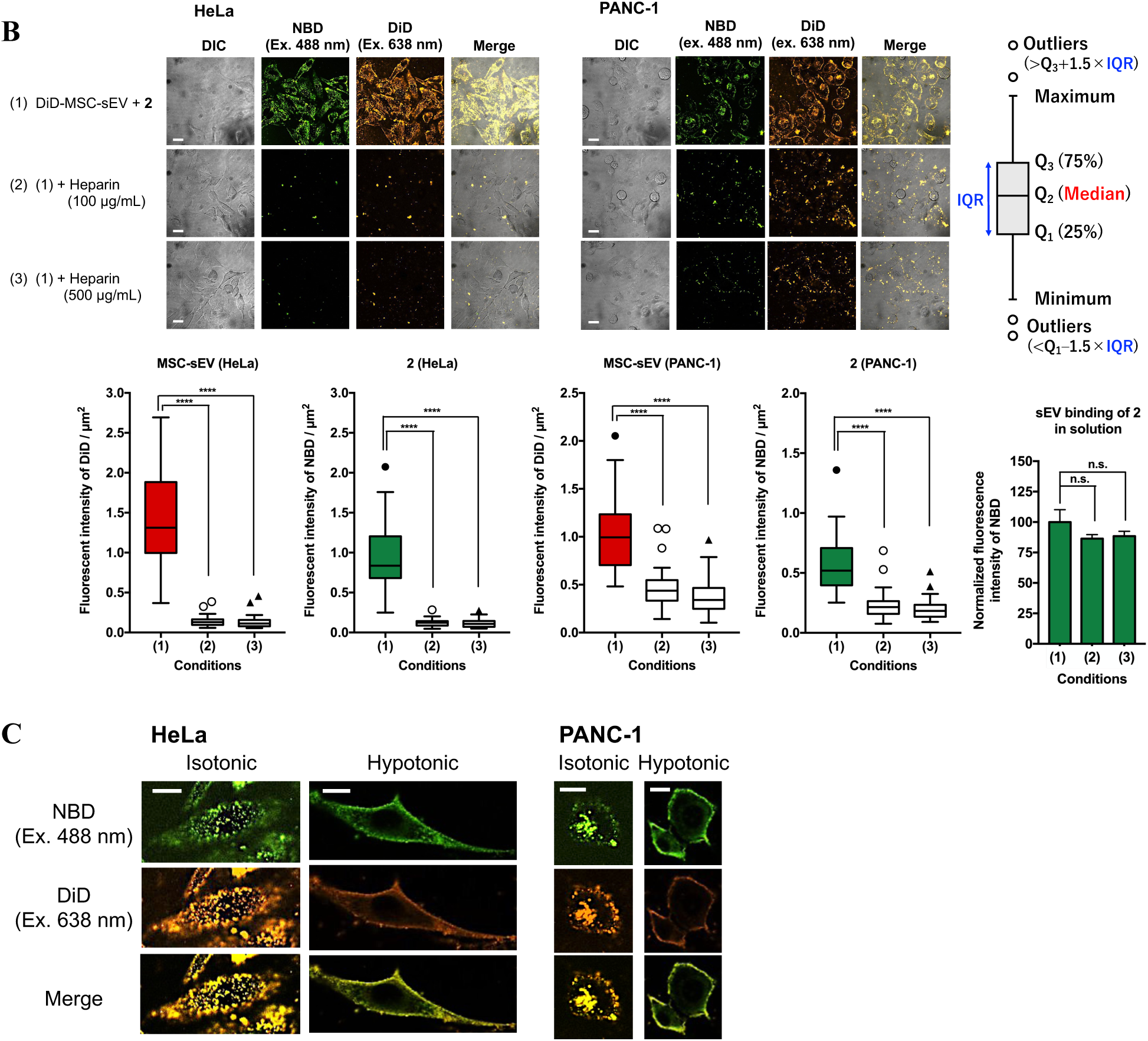
Elucidation of the mechanism by which **2** facilitates the accumulation of sEVs on the cell surface and its subsequent endocytosis. (A) Schematic diagram of the proposed mechanism: (*i*) **2**-modified sEVs accumulate on the cell surface *via* electrostatic interactions between positively charged **2** on the sEVs and negatively charged molecules, such as heparan sulfates of membrane proteins and phosphatidylserine (PS); (*ii*) **2**-modified sEVs are internalized into cells following deformation of the plasma membrane. The inhibition points of heparin and hypotonicity are shown in this panel. (B) Cellular uptake of DiD-MSC-sEVs and **2** under conditions (1) 0, (2) 100, and (3) 500 µg/mL heparin after incubation for more than 1 h. The scale bar is 10 µm. Fluorescence intensities of DiD and NBD per cell area (µm^2^), analyzed from the confocal images, are shown in box-and-whisker plots (mean ± S.D., *n* = 40 cells). sEV binding of **2** was also evaluated in solution in the presence of heparin (mean ± S.D., *n* = 3). Conditions (1)–(3) correspond to those shown in the confocal images. Statistical analysis was performed using Student’s *t*-test at an alpha level of 0.05 (n.s.: not significant, \*\*\*\**p* < 0.0001). (C) Internalization of DiD-MSC-sEVs and **2** under isotonic or hypotonic conditions after incubation for more than 1 h. The scale bar is 5 µm.

Although we designed derivatives of **2** (entries **4**–**7**) (Table 1 and Figures S4–S7) to further improve the cellular uptake of peptides, peptide **2** was found to be the most effective functional peptide. Peptide **4** has a structure similar to that of **2** and was used as a control for peptides **5**–**7**. To enable consistent evaluation, peptide designs **4**–**7** were unified into an NBD-branched form. All eight lysine residues of peptide **5** were replaced with arginine residues because arginine-rich peptides interact more strongly with anionic membranes than lysine-rich peptides^37, 38^. Peptides **6** and **7** have a branched NBD-labeling form with octanoic acid introduced to the main chain, as modification with hydrocarbon chains has been reported to facilitate peptidic interaction with membranes^39, 40^. In the presence of excess liposomes (20 µM), the peptide uptake ratio with and without liposomes was used as the evaluation criterion. Consequently, peptide **4** exhibited the most effective cellular uptake in the two cancer cell lines but not in vascular endothelial cells (HUVECs), with negligible intracellular signals (Figures S11–S13), indicating that **4** is a preferable peptide that is efficiently transported into cells in the presence of vesicles but is not taken up by cells alone. Arginine substitution and hydrocarbon chain modification were not suitable for this study.

### Identification of endocytosis pathways for sEV uptake in the presence of an sEV-modifying peptide

We found that **2** enhances the endocytic uptake of DiD-MSC-sEVs through a clathrin-mediated pathway in two types of cancer cells, which is a completely different phenomenon from that of DiD-MSC-sEVs alone. To determine the endocytic uptake pathways of DiD-MSC-sEVs, we investigated the reduction in cellular uptake with various endocytosis inhibitors. EIPA (inhibition of the Na⁺/H⁺ exchanger)^41^, wortmannin (inhibition of PI3K activity)^42^, and cytochalasin D (inhibition of actin polymerization)^43^ were used as macropinocytosis inhibitors. Wortmannin and cytochalasin D are also potentially effective in inhibiting phagocytosis^42, 43^. Pitstop2 (associated with the clathrin terminal domain)^44^ and dynasore (inhibition of dynamin GTPase activity)^45^ were used as clathrin-mediated endocytosis inhibitors. Nystatin (associated with sterol lipids)^46^ is an inhibitor of caveolae-mediated endocytosis. In the presence of **2**, the uptake of DiD-MSC-sEVs decreased remarkably in HeLa and PANC-1 cells treated with pitstop2 [condition (6)] (Figure 4). To confirm the reproducibility of DiD-MSC-sEV uptake *via* the clathrin-mediated endocytic pathway, we treated cells with pitstop2 and dynasore, which have different modes of action [conditions (6)(7)]. Consequently, the uptake of DiD-MSC-sEVs decreased to a level similar to that in untreated cells [condition (1)] (Figure 4). Both NBD and DiD fluorescence signals were reduced in the presence of these inhibitors, indicating that the endocytic uptake of **2**-modified sEVs was highly dependent on the clathrin-mediated pathway (Figure 4). Macropinocytosis (or phagocytosis) [conditions (3)–(5)] and caveolae-mediated endocytosis (data not shown) were not involved in the uptake. By contrast, the uptake of DiD-MSC-sEVs at the same concentration (1 × 10^8^ particles/mL) was poorly observed in the absence of **2** (Figure 5A); therefore, a 10-fold higher concentration (1 × 10^9^ particles/mL) was used to image the uptake of DiD-MSC-sEVs alone (Figure S14). The cellular uptake of sEVs alone decreased slightly when the cells were treated with pitstop2 and dynasore (Figure S14), but their uptake may not follow a single pathway (Figure 4), as various endocytic pathways have been reported for sEVs in previous studies^47^.

**Figure 4.**
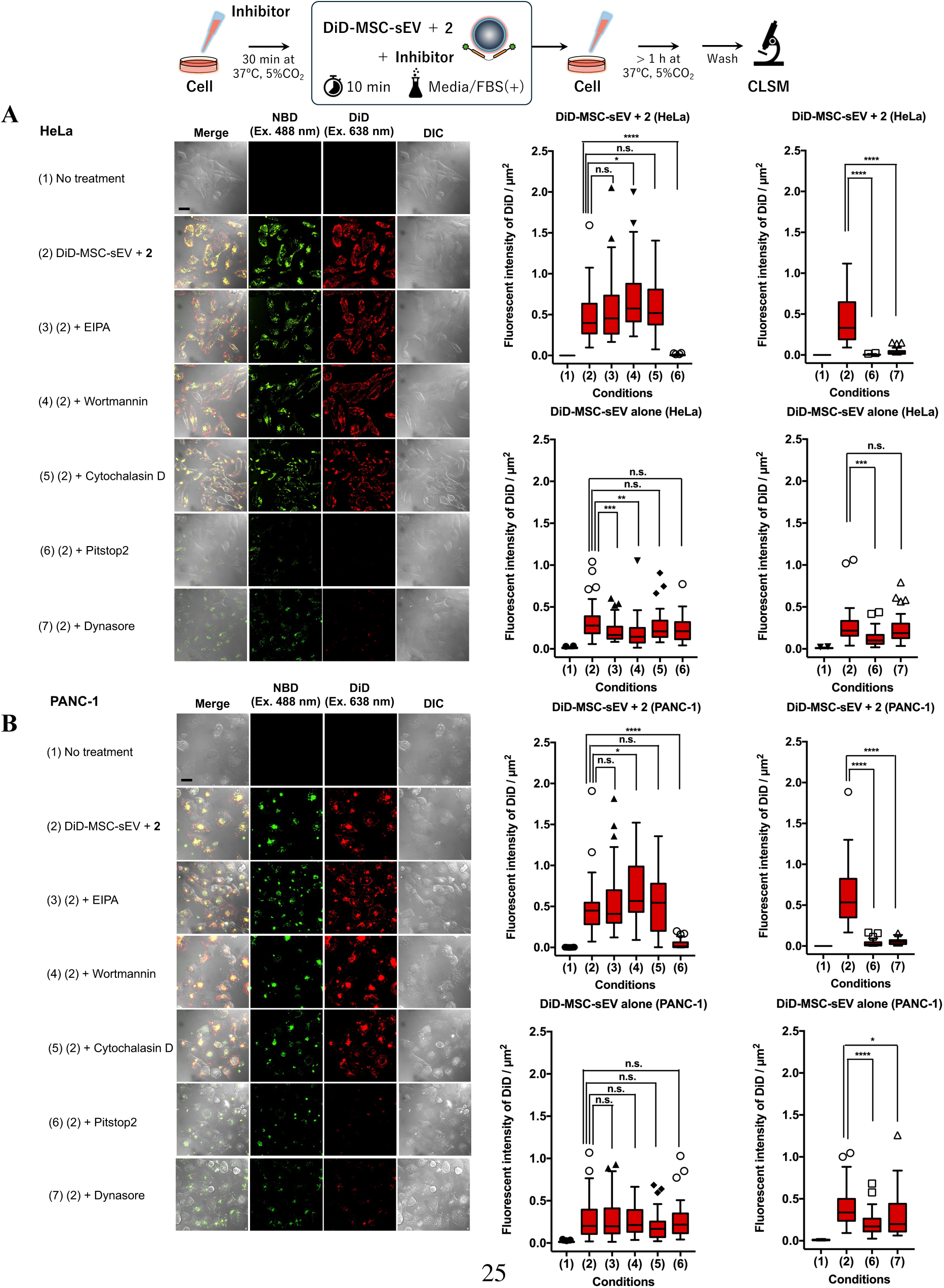
Inhibition of sEV endocytosis. In the presence of **2**, DiD-MSC-sEV uptake was observed at 1 × 10^8^ particles/mL in (A) HeLa and (B) PANC-1 cells treated with endocytosis inhibitors (10 µM EIPA, 1 µM wortmannin, 200 nM cytochalasin D, 15 µM pitstop2, or 15 µM dynasore). In the absence of **2**, DiD-MSC-sEV uptake was observed at 1 × 10^9^ particles/mL. The scale bar is 10 µm. Fluorescence intensities of DiD per cell area (µm^2^) were analyzed from the confocal images (mean ± S.D., *n* = 40 cells) (refer to Figure S14). Statistical analysis was performed using Student’s *t*-test at an alpha level of 0.05 (n.s.: not significant, \**p* < 0.05, \*\**p* < 0.01, \*\*\**p* < 0.001, \*\*\*\**p* < 0.0001).

### Enhanced cellular uptake of sEVs and its application for a drug delivery system using an sEV-modifying peptide

We found that the cellular uptake of DiD-MSC-sEVs and **2** under coexisting conditions increased to over fivefold and 20-fold higher, respectively, than that of each alone, and that the sEV-modifying peptide is useful for easily loading anticancer drugs onto the sEV surface and delivering them into cancer cells. To evaluate cellular uptake of sEVs in the presence of **2**, two types of cancer cells were treated with DiD-MSC-sEVs (6 × 10^7^ particles/mL) with or without **2** at 37 °C for more than 1 h. In the absence of **2**, DiD fluorescence was observed in only a limited number of cells, whereas in the presence of **2**, the fluorescence intensity increased in almost all cells (Figure 5A). Interestingly, in the absence of DiD-MSC-sEVs, NBD signals were negligible, similar to those in untreated cells. Data analysis supported these findings; a positive correlation between NBD and DiD fluorescence intensities per cell area was observed for both cell lines only under coexisting conditions [Figure 5B, condition (4)]. Notably, although DiD-MSC-sEVs and **2** exhibited high colocalization coefficients when accumulated on the cell surface (Figure 2E), their correlation factor values (*R*) inside the cells were 0.71 and 0.65 for HeLa and PANC-1 cells, respectively (Figure 5B), indicating that **2** begins to dissociate from DiD-MSC-sEVs inside the cells under reducing conditions, facilitating conversion from the dimeric to the monomeric form (Figure 2A). The mean DiD fluorescence values of DiD-MSC-sEVs in the presence of **2** were 5.1- and 5.7-fold higher than those in the absence of **2** in HeLa and PANC-1 cells, respectively [Figure 5C; conditions (2) vs. (4)]. Similarly, the NBD fluorescence of **2** in the presence of DiD-MSC-sEVs was 24.6- and 21.8-fold higher than that in the absence of DiD-MSC-sEVs in HeLa and PANC-1 cells, respectively [Figure 5C; conditions (3) vs. (4)].

**Figure 5.**
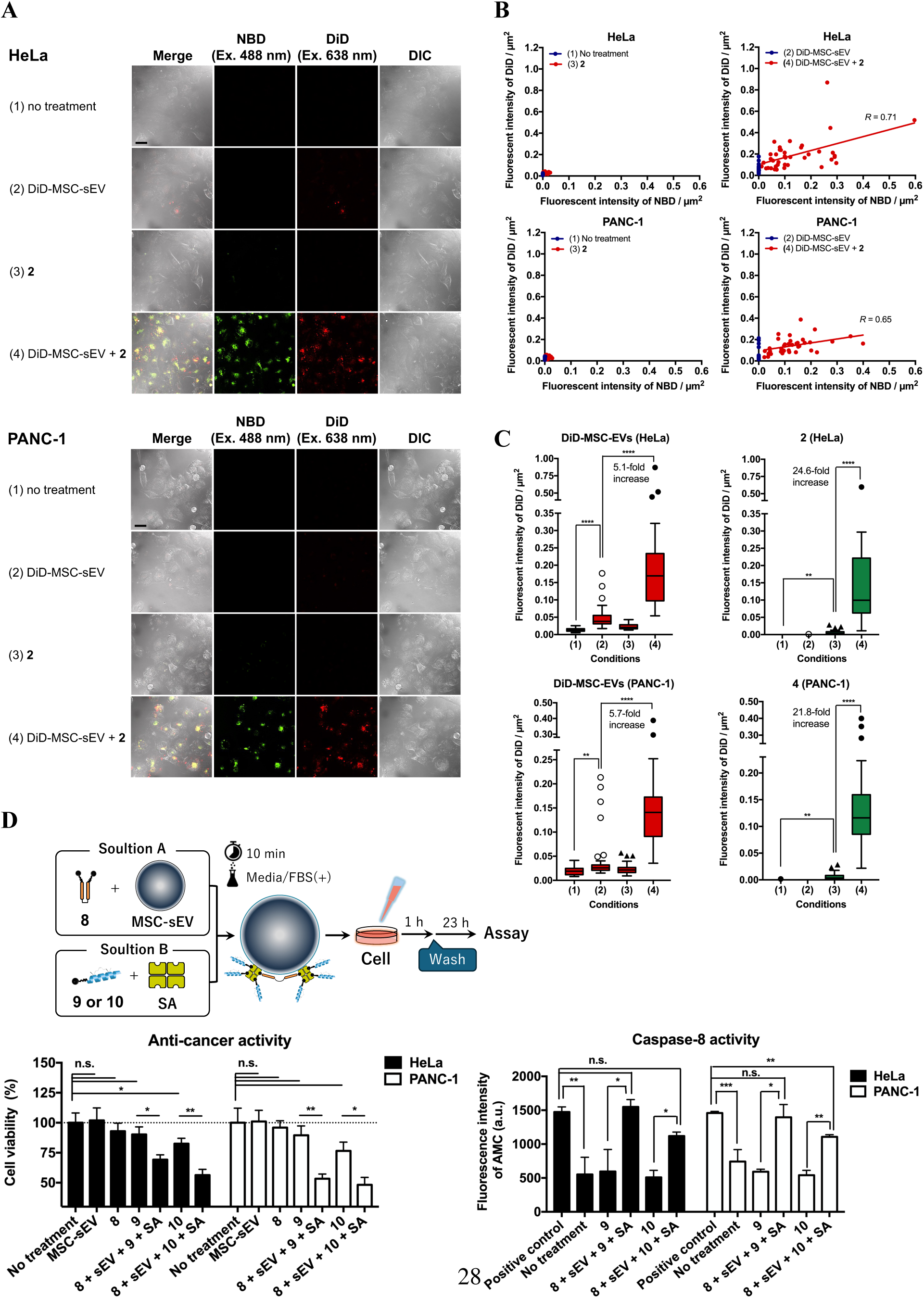
Enhanced cellular uptake of both DiD-MSC-sEVs and **2**, and application to a drug delivery system using anticancer peptides. (A) Observation of cellular uptake of DiD-MSC-sEVs alone, **2** alone, and their mixture (sEVs: 6 × 10^7^ particles/mL; **2**: 1 µM). The scale bar is 10 µm. (B) Correlation of cellular uptake between DiD-MSC-sEVs and **2**. Fluorescence intensity of DiD per cell area (µm^2^), as measured from confocal images, was plotted against that of NBD (*n* = 40 cells). Correlation coefficients (*R*) were calculated as 0.71 and 0.65 for HeLa and PANC-1 cells, respectively, under condition (4). (C) Comparison of cellular uptake of DiD-MSC-sEVs (red) and **2** (green) (mean ± S.D., *n* = 40 cells). The median fluorescence intensities of DiD/µm^2^ in the presence of **2** were increased 5.1- and 5.7-fold compared with those in the absence of **2** for HeLa and PANC-1 cells, respectively [condition (2) vs. (4)]. The median fluorescence intensities of NBD/µm^2^ in the presence of DiD-MSC-sEVs were increased 24.6- and 21.8-fold compared with those in the absence of DiD-MSC-sEVs for HeLa and PANC-1 cells, respectively [condition (3) vs. (4)]. Statistical analysis was performed using Student’s *t*-test at an alpha level of 0.05 (\*\**p* < 0.01, \*\*\*\**p* < 0.0001). (D) Biotinylated di-nFAAV5 (**8**) and membrane-impermeable stapled anticancer peptides (**9** and **10**) were loaded onto the surface of non-labeled MSC-sEVs *via* streptavidin (SA). Cell viabilities and caspase-8 activities of HeLa and PANC-1 cells were evaluated under the same conditions (mean ± S.D., *n* = 3). Statistical analysis was performed using Student’s *t*-test at an alpha level of 0.05 (n.s.: not significant, \**p* < 0.05, \*\**p* < 0.01, \*\*\**p* < 0.001).

We developed a simple drug-loading system for sEVs using biotinylated sEV-modifying peptides and drugs that form a complex *via* SA on the sEV surface, as illustrated in Figure 5D. Accordingly, we prepared N-terminus-biotinylated di-nFAAV5 (entry **8**) and anticancer stapled peptides biotin-SAH-p53-4 (entry **9**)^48^ and biotin-BIM-BH3 (entry **10**)^49^ (Table 1 and Figures S8– S10), which are membrane-impermeable and apoptosis-inducing peptides, respectively. No cytotoxicity of **8** was observed in HeLa and PANC-1 cells under either condition; **8** was delivered into cells using nontoxic liposomes (Figure S15) and existed alone in solution (Figure 5D). By contrast, the anticancer activities of **9** and **10** were clearly demonstrated using this drug-loading system on non-labeled MSC-sEVs; namely, a remarkable decrease in cell viability and increase in caspase-8 activity were observed for **9** and **10** in the presence of **8** and MSC-sEVs (Figure 5D). No membrane perturbation was observed under the same conditions; weak signals from the membrane-impermeable dye PI were detected in only 2–7% of the cells (Figures S16 and S17). Collectively, these data suggest that the sEV-modifying peptide, using MSC-sEVs as a drug carrier, successfully delivered anticancer drugs into cancer cells and induced apoptosis. The simplicity and efficiency of this sEV-based drug-loading system are key advantages for drug delivery applications.

## Discussion

In the present study, we found that enhanced cellular uptake of sEVs and **2** was observed only under coexisting conditions, whereas limited uptake of sEVs and **2** was observed under single conditions (Figure 5A–C). This suggests that sEVs and **2** require each other for effective cellular uptake, and **2** utilizes sEVs as a scaffold to exert its function by accumulating highly on the sEV surface.

The mechanism underlying the enhanced cellular uptake of sEVs in the presence of the sEV-modifying peptide **2** is shown in Figure 6. In the first step (Figure 6), the electrostatic interaction between **2** bound to sEVs and the cell membrane acts as a driving force to enhance the accumulation of sEVs on the cell surface (Figure 3B). The cellular uptake of DiD-MSC-sEVs in the absence and presence of **2** was remarkably different (Figure 5A), even though their ζ-potential values were similar (Figure 2D). This can be explained by the structural differences between DiD and **2**; that is, the lipophilic carbocyanine dye DiD deeply inserts into the membrane^50^ so that the positively charged nitrogen atom may be buried at a position unsuitable for interaction with negatively charged molecules on the plasma membrane (Figure 6). By contrast, the dimeric curvature-sensing peptide adopts an amphipathic α-helix structure (Figures 1C and 2B), with eight Lys residues located on the hydrophilic side, facing the aqueous environment and suitable for interaction with negatively charged molecules on the cell surface (Figure 6). We clearly demonstrated that enhanced cellular accumulation of sEVs is achieved by the function of **2** and does not depend on either the ζ-potential or DiD labeling of sEVs (Figure 2E and F). In the second step (Figure 6), negative membrane curvature prior to endocytic uptake was induced by sEV accumulation on the cell surface (Figure 3C). Although the driving force that initiates endocytosis remains unknown, three factors plausibly contribute to this phenomenon: (*i*) **2**, which accumulates highly on sEVs, induces plasma membrane deformation to fit the shape of sEVs; (*ii*) mechanosensitive ion channels on cancer cells deform the cell membrane in response to mechanical stimuli^51^; and (*iii*) the proximity of anionic molecules, such as PS^27^, which are recruited through interaction with **2** on sEVs, induces membrane curvature. In the third step (Figure 6), **2**-modified sEVs were internalized *via* the clathrin-mediated endocytosis pathway (Figure 4). These results were unexpected but highly reliable, as sEV internalization was completely inhibited using multiple inhibitors with different modes of action against clathrin-mediated endocytosis. Clathrin is a self-assembling protein that prefers curved membranes^52^ and spontaneously coats curved membranes from the cytoplasmic side to form clathrin-coated pits for endocytosis. Endocytosis induced by **2**-modified sEVs was distinct from that induced by untreated sEVs (Figure 4). Although various endocytosis pathways, such as clathrin-independent or caveolae-mediated pathways, have been reported^11, 47^, sEV-modifying peptides can alter the cellular uptake pathway of sEVs. We ruled out the involvement of receptor-mediated endocytosis in the internalization of **2**-modified sEVs because if **2** were recognized by unknown receptors, internalization of **2** alone would be observed in the same manner.

**Figure 6.**
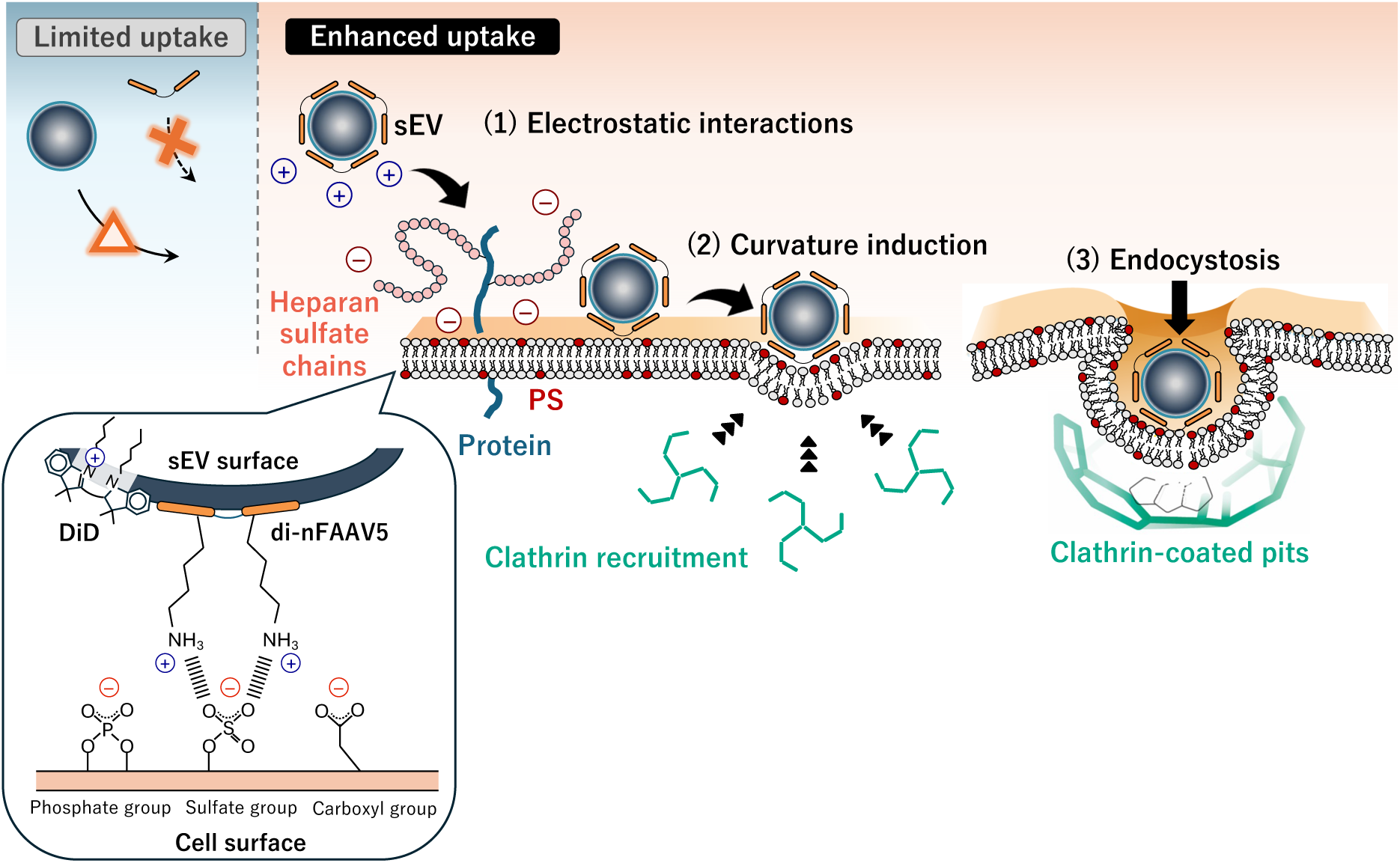
Proposed model of an sEV-modifying peptide facilitating cellular uptake of sEVs. Step (1): Electrostatic interactions between **2** on sEVs and negatively charged molecules (including heparan sulfate chains and PS) on the plasma membrane serve as a driving force for sEV accumulation on the cell surface. Detailed interactions are shown in an enlarged view. Step (2): The **2**-modified sEV surface induces deformation of the plasma membrane, followed by the recruitment of clathrin from the cytoplasmic side. Clathrin, a curvature-sensing/generating protein, forms clathrin-coated pits. Step (3): sEVs coated with **2** are internalized *via* the clathrin-mediated endocytosis pathway.

Previously, it was reported that EV labeling methods can influence the properties and fate of EVs, such as their interaction with recipient cells and modulation of their activity, and that incorporation of reporter proteins into EVs is desirable^53^. However, it is difficult to achieve high expression levels of reporter proteins in EVs, and low drug-loading efficiency remains an issue. Moreover, since some types of EV-sorting proteins are heterogeneously loaded onto EVs^54^, the target proteins used for cargo loading must be carefully considered. If a drug delivery system using EVs can be artificially controlled, altering the inherent properties of EVs is not a major concern; therefore, EV labeling methods should be chosen based on the research objective.

Anticancer peptides **9** and **10** were compatible with this simple drug-loading system onto sEVs and induced apoptosis in two types of cancer cells, as they did not exhibit strong direct membrane penetration because of their hydrophilicity (Figure 5D). Some of the loaded drugs likely escaped from endosomes after membrane rupture triggered by the proton sponge effect^55^ and exerted anticancer activity after being transferred to the cytosol (Figure 5D). We also attempted to load a hydrophobic compound (camptothecin) onto sEVs, which induces apoptosis in cancer cells *via* specific inhibition of DNA topoisomerase I^56^. However, this drug-loading system did not work effectively for camptothecin because of its high hydrophobicity (data not shown). These results indicate that if the anticancer drug is too hydrophobic, the function and properties of the system will be strongly affected by the loaded drug. The anticancer activity of MSC-sEVs alone was observed in PBS(+), but not in DMEM(+) (Figure S15B), indicating that their activity at low particle concentrations depends on the condition.

In future studies, the next generation of sEV-modifying peptides enabling covalent binding to sEVs would be a promising tool for drug delivery *in vivo*, as covalent bonding prevents the peptide from switching binding partners, even in the presence of a mixture of different sEVs. The advantage of this system is that multiple functional units can be loaded onto the sEV surface simultaneously using the sEV-modifying peptide, making it possible to achieve controlled drug delivery and release to target specific tissues and track the biokinetics of the sEVs by loading dyes onto the sEV surface. Curvature-sensing peptides have been proven to show protease resistance against trypsin and proteinase K^57^, and **2**-modified sEVs are taken up by cancer cells even at low particle concentrations but are poorly taken up by HUVECs, making these properties potentially suitable for *in vivo* study applications.

## Conclusion

The sEV-modifying peptide exerted its functions especially at low particle concentrations by highly enhancing the accumulation of sEVs on the cell surface and their endocytic uptake *via* a clathrin-mediated pathway. Cellular uptake of sEVs and the sEV-modifying peptide was facilitated in cancer cells by over fivefold and 20-fold, respectively, under coexisting conditions compared with those administered alone. The sEV-modifying peptide functions as a *membrane interfactant* to reduce the energy barrier for the cellular uptake of sEVs. Moreover, the sEV-modifying peptide allowed us to load anticancer drugs onto the surface of sEVs and induce apoptosis in cancer cells.

## ASSOCIATED CONTENT

### Supporting Information

The Supporting Information is available free of charge at https://XXX.

HPLC profiles and MS analyses of **1**–**10**, cellular uptake of ATTO647N-liposomes (HeLa, PANC-1, and HUVEC), cellular uptake of DiD-MSC-sEVs in the presence of inhibitors, cytotoxicity of peptides and anti-cancer activity of MSC-sEVs, and membrane perturbation assay (HeLa and PANC-1) (PDF).

### Author Contributions

K.K.: Conceptualization, Resources, Funding acquisition, Project administration, Investigation, Formal analysis, Methodology, Writing – original draft, Writing – review and editing, Visualization, Supervision. K.H.: Investigation, Formal analysis. A.T.: Formal analysis. Y.O.: Investigation. Y.K.: Investigation. K.M.: Resources, Writing – review and editing, Supervision.

### Notes

The authors declare no competing financial interest.

## ACKNOWLEDGMENT

This study was financially supported by the Chubei Itoh Foundation (K.K.) and JSPS KAKENHI (21K15254 to K.K.). The authors thank Prof. Dr. Yukio Kato (Hiroshima University) for providing the MSC-R37 cells.

## ABBREVIATIONS

ATTO647N-DPPE: 1,2-dipalmitoyl-*sn*-glycero-3-phosphoethanolamine labeled with ATTO 647N;
bFGF: basic fibroblast growth factor;
CD: circular dichroism;
CLSM: confocal laser scanning microscopy;
di-nFAAV5: dimeric nFAAV5;
DiD: 1,1′-dioctadecyl-3,3,3′,3′-tetramethylindodicarbocyanine;
DMEM: Dulbecco’s modified Eagle’s medium;
DMF: *N,N*-dimethylformamide;
EV-CaRiS: extracellular vesicle catch-and-release isolation system;
FBS: fetal bovine serum;
HUVEC: vascular endothelial cells;
MSC: mesenchymal stem cell;
NBD: nitrobenzoxadiazole;
PBS: phosphate-buffered saline;
PI: propidium iodide;
POPC: 1-palmitoyl-2-oleoyl-*sn*-glycero-3-phosphocholine;
POPS: 1-palmitoyl-2-oleoyl-*sn*-glycero-3-phospho-L-serine;
PS: phosphatidylserine;
SA: streptavidin;
sEV: small extracellular vesicles.

## Table of Contents

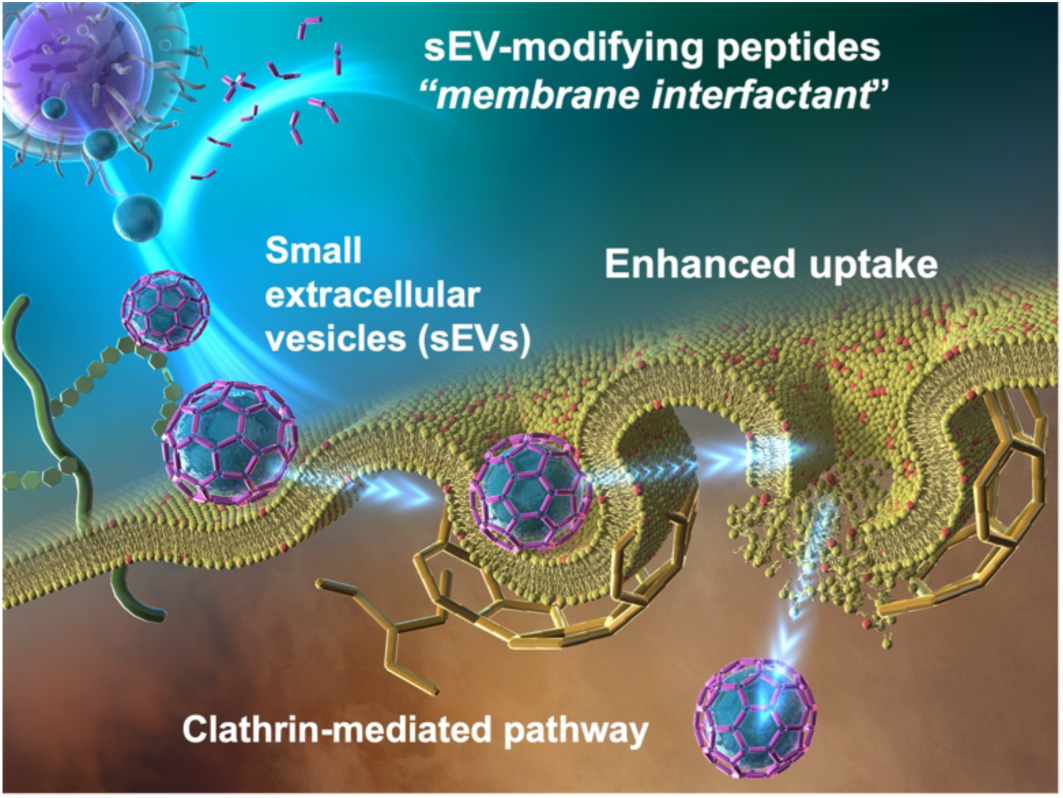

