## Supplemental Information for "Curvature-sensing peptide functions as a membrane interfactant that glues small extracellular vesicles to cell membranes and enhances vesicle cellular uptake"

### Table of Contents

**Supplementary Figures**


| | | |
|---|---|---|
| Figure S1 | HPLC profile and MS analysis of **1** (NBD-nFAAV5) | P. 3 |
| Figure S2 | HPLC profile and MS analysis of **2** (di-NBD-nFAAV5) and **2'** (di-nFAAV5) | P. 4–5 |
| Figure S3 | HPLC profile and MS analysis of **3** (di-nFAAV5-NBD) | P. 6 |
| Figure S4 | HPLC profile and MS analysis of **4** [di-K(NBD)-nFAAV5] | P. 7 |
| Figure S5 | HPLC profile and MS analysis of **5** [di-K(NBD)-nFAAV6] | P. 8 |
| Figure S6 | HPLC profile and MS analysis of **6** [di-C8-K(NBD)-nFAAV5] | P. 9 |
| Figure S7 | HPLC profile and MS analysis of **7** [di-C8-K(NBD)-nFAAV6] | P. 10 |
| Figure S8 | HPLC profile and MS analysis of **8** (di-biotin-nFAAV5) | P. 11 |
| Figure S9 | HPLC profile and MS analysis of **9** (biotin-SAH-p53-4) | P. 12 |
| Figure S10 | HPLC profile and MS analysis of **10** (biotin-BIM BH3) | P. 13 |
| Figure S11 | Cellular uptake of ATTO647N-liposomes (HeLa) | P. 14–15 |
| Figure S12 | Cellular uptake of ATTO647N-liposomes (PANC-1) | P. 16 |
| Figure S13 | Cellular uptake of ATTO647N-liposomes (HUVEC) | P. 17 |
| Figure S14 | Cellular uptake of DiD-MSC-sEVs in the presence of inhibitors | P. 18 |
| Figure S15 | Cytotoxicity of peptides and anti-cancer activity of MSC-sEVs | P. 19 |
| Figure S16 | Membrane perturbation assay (HeLa) | P. 20 |
| Figure S17 | Membrane perturbation assay (PANC-1) | P. 21 |

**1** (NBD-nFAAV5)

**NBD-DKBLLKXLNKBTDBLSKX-GSGSK-$NH_2$**

**A**

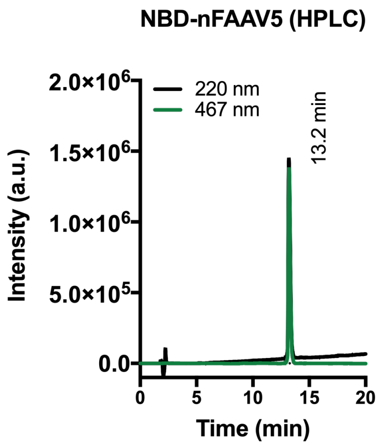


**B** Calculated values

$[M+H]^+$ = 2575.46

$[M+2H]^{2+}$ = 1288.23

$[M+3H]^{3+}$ = 859.16

$[M+4H]^{4+}$ = 644.62

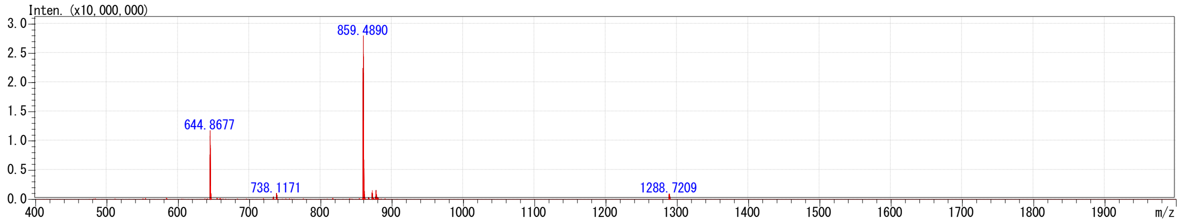


**Figure S1. HPLC profile and MS analysis of 1 (NBD-nFAAV5).**

The HPLC profile (A) and the ESI-MS analytical data (B) of the purified peptide. (A) The purity and the retention time were analyzed by RP-HPLC on a COSMOSIL $5C_{18}$-AR-II column (4.6 mm I.D. × 150 mm) using a linear gradient from 30 to 80% acetonitrile in 0.1% aqueous TFA for 20 min at 40 ºC at a flow rate of 1.0 mL/min. The purity of the peptide was calculated to be over 95% based on the peak area. (B) The calculated masses of multivalent ions are described on top of the observed *m/z* data.

**2** (di-NBD-nFAAV5)

**NBD-DKBLLKXLNKBTDBLSKX-GSGS-C-$NH_2$**

**NBD-DKBLLKXLNKBTDBLSKX-GSGS-C-$NH_2$**

**A** **di-NBD-nFAAV5 (HPLC)**

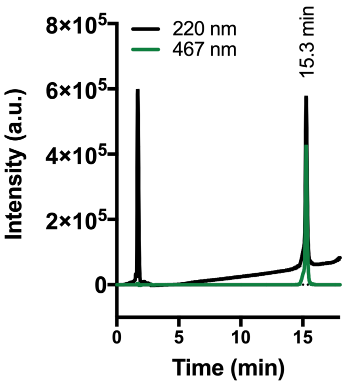


Calculated values of **2**

$[M+H]^{+}$ = 5097.72

$[M+2H]^{2+}$ = 2549.36

$[M+3H]^{3+}$ = 1699.91

$[M+4H]^{4+}$ = 1275.18

$[M+5H]^{5+}$ = 1020.34

$[M+6H]^{6+}$ = 850.45

**B**

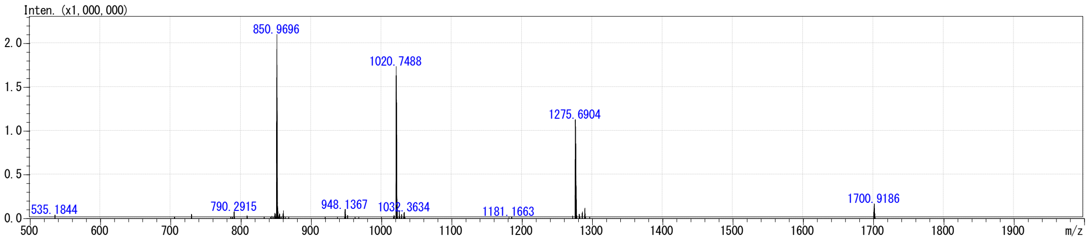


**2'** (di-nFAAV5 used for measuring CD and Z-potential)

**DKBLLKXLNKBTDBLSKX-GSGS-C-$NH_2$**

**DKBLLKXLNKBTDBLSKX-GSGS-C-$NH_2$**

**C** **di-nFAAV5 (HPLC)**

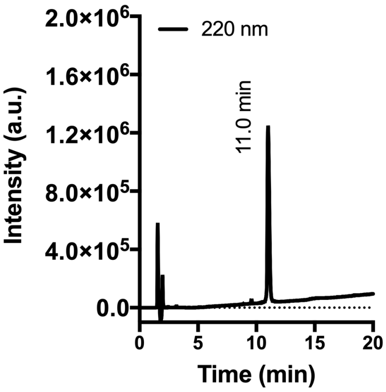


Calculated values of **2'**

$[M+H]^{+}$ = 4771.72

$[M+2H]^{2+}$ = 2386.36

$[M+3H]^{3+}$ = 1591.24

$[M+4H]^{4+}$ = 1193.68

$[M+5H]^{5+}$ = 955.14

$[M+6H]^{6+}$ = 796.12

**D**

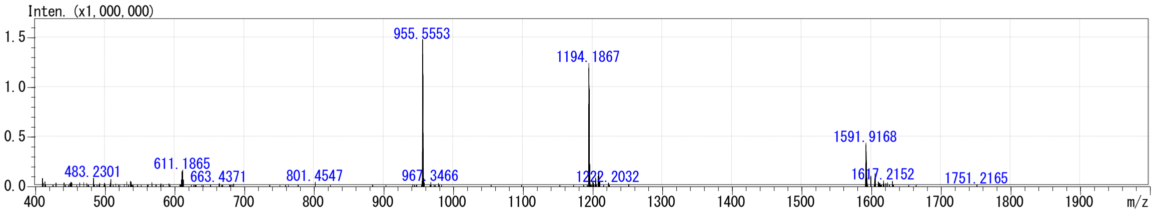

**Figure S2. HPLC profile and MS analysis of 2 (di-NBD-nFAAV5) and 2' (di-nFAAV5).**

The HPLC profiles (A)(C) and the ESI-MS analytical data (B)(D) of the purified peptides. (A)(C) The purities and the retention times were analyzed by RP-HPLC on a COSMOSIL 5$C_{18}$-AR-II column (4.6 mm I.D. × 150 mm) using a linear gradient from 30 to 80% acetonitrile in 0.1% aqueous TFA for 20 min at 40 ºC at a flow rate of 1.0 mL/min. The purities of the peptides was calculated to be over 95% based on the peak area. (B)(D) The calculated masses of multivalent ions are described on top of the observed *m/z* data.

**3** (di-nFAAV5-NBD)

**DKBLLKXLNKBTDBLSKX-GSGSK(NBD)-C-$NH_2$**

**|**

**DKBLLKXLNKBTDBLSKX-GSGSK(NBD)-C-$NH_2$**

**A**

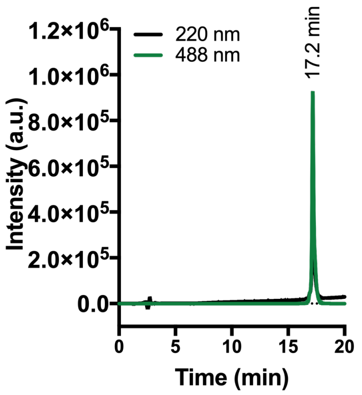


**B**

Calculated values

$[M+H]^+$ = 5353.91

$[M+4H]^{4+}$ = 1339.23

$[M+5H]^{5+}$ = 1071.58

$[M+6H]^{6+}$ = 893.15

$[M+7H]^{7+}$ = 765.70

$[M+8H]^{8+}$ = 670.11

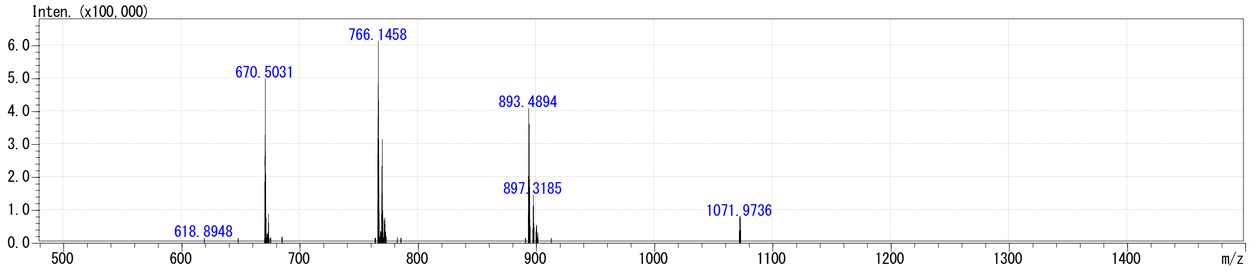


**Figure S3. HPLC profile and MS analysis of 3 (di-nFAAV5-NBD).**

The HPLC profile (A) and the ESI-MS analytical data (B) of the purified peptide. (A) The purity and the retention time were analyzed by RP-HPLC on a COSMOSIL $5C_{18}$-AR-II column (4.6 mm I.D. × 150 mm) using a linear gradient from 30 to 80% acetonitrile in 0.1% aqueous TFA for 20 min at 40 ºC at a flow rate of 1.0 mL/min. The purity of the peptide was calculated to be over 95% based on the peak area. (B) The calculated masses of multivalent ions are described on top of the observed *m/z* data.

**4** [di-K(NBD)-nFAAV5]

**K(NBD)-DKBLLKXLNKBTDBLSKX-GSGS-CK-NH$_2$**

**K(NBD)-DKBLLKXLNKBTDBLSKX-GSGS-CK-NH$_2$**

**A**

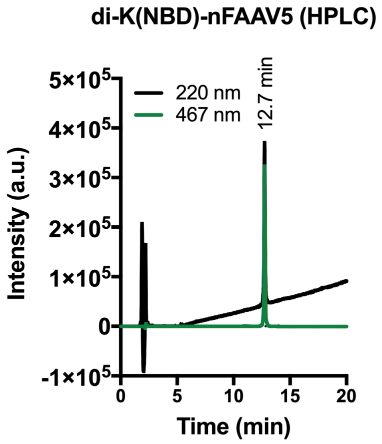


**B**

Calculated values

$[M+H]^+$ = 5610.10

$[M+4H]^{3+}$ = 1870.70

$[M+4H]^{4+}$ = 1403.28

$[M+5H]^{5+}$ = 1122.83

$[M+6H]^{6+}$ = 935.86

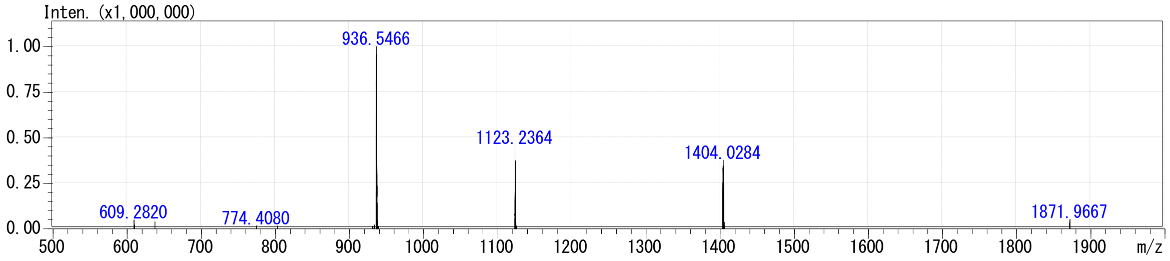


**Figure S4. HPLC profile and MS analysis of 4 [di-K(NBD)-nFAAV5].**

The HPLC profile (A) and the ESI-MS analytical data (B) of the purified peptide. (A) The purity and the retention time were analyzed by RP-HPLC on an InterSustainSwift 5C$_{18}$-AR-II column (4.6 mm I.D. × 150 mm) using a linear gradient from 30 to 80% acetonitrile in 0.1% aqueous TFA for 20 min at 40 ºC at a flow rate of 1.0 mL/min. The purity of the peptide was calculated to be over 95% based on the peak area. (B) The calculated masses of multivalent ions are described on top of the observed *m/z* data.

**5** [di-K(NBD)-nFAAV6]

**K(NBD)-DRBLLRXLNRBTDBLSRX-GSGS-CK-NH$_2$**

**K(NBD)-DRBLLRXLNRBTDBLSRX-GSGS-CK-NH$_2$**

**A**

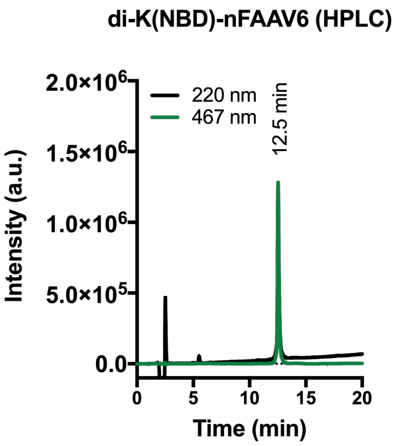


**B**

Calculated values

$[M+H]^+ = 5834.15$

$[M+4H]^{4+} = 1459.29$

$[M+5H]^{5+} = 1167.64$

$[M+6H]^{6+} = 973.20$

$[M+7H]^{7+} = 834.31$

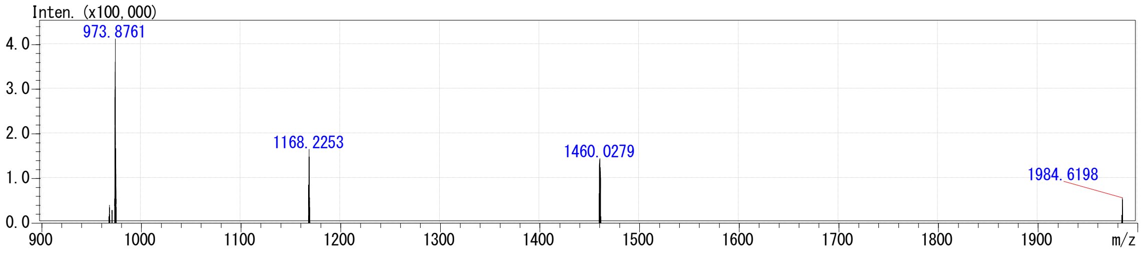


**Figure S5. HPLC profile and MS analysis of 5 [di-K(NBD)-nFAAV6].**

The HPLC profile (A) and the ESI-MS analytical data (B) of the purified peptide. (A) The purity and the retention time were analyzed by RP-HPLC on an InterSustainSwift 5C$_{18}$-AR-II column (4.6 mm I.D. × 150 mm) using a linear gradient from 30 to 80% acetonitrile in 0.1% aqueous TFA for 20 min at 40 ºC at a flow rate of 1.0 mL/min. The purity of the peptide was calculated to be over 95% based on the peak area. (B) The calculated masses of multivalent ions are described on top of the observed *m/z* data.

**6** [di-C8-K(NBD)-nFAAV5]

**C8-K(NBD)-DKBLLKXLNKBTDBLSKX-GSGS-CK-NH$_2$**

**|**

**C8-K(NBD)-DKBLLKXLNKBTDBLSKX-GSGS-CK-NH$_2$**

**A**

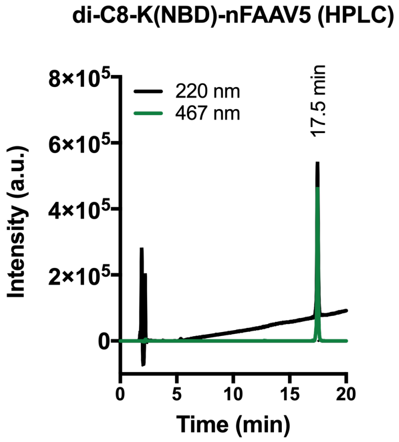


**B**

Calculated values

$[M+H]^+ = 5862.31$

$[M+4H]^{4+} = 1466.33$

$[M+5H]^{5+} = 1173.27$

$[M+6H]^{6+} = 977.89$

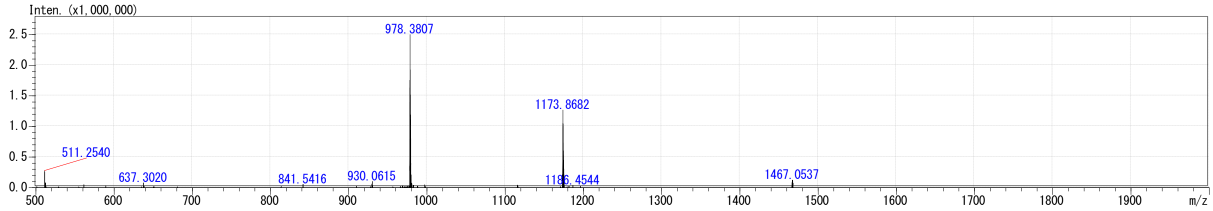


**Figure S6. HPLC profile and MS analysis of 6 [di-C8-K(NBD)-nFAAV5].**

The HPLC profile (A) and the ESI-MS analytical data (B) of the purified peptide. (A) The purity and the retention time were analyzed by RP-HPLC on an InterSustainSwift 5C$_{18}$-AR-II column (4.6 mm I.D. × 150 mm) using a linear gradient from 30 to 80% acetonitrile in 0.1% aqueous TFA for 20 min at 40 ºC at a flow rate of 1.0 mL/min. The purity of the peptide was calculated to be over 95% based on the peak area. (B) The calculated masses of multivalent ions are described on top of the observed *m/z* data.

**7** [di-C8-K(NBD)-nFAAV6]

**C8-K(NBD)-DRBLLRXLNRBTDBLSRX-GSGS-CK-$NH_2$**

**C8-K(NBD)-DRBLLRXLNRBTDBLSRX-GSGS-CK-$NH_2$**

**A**

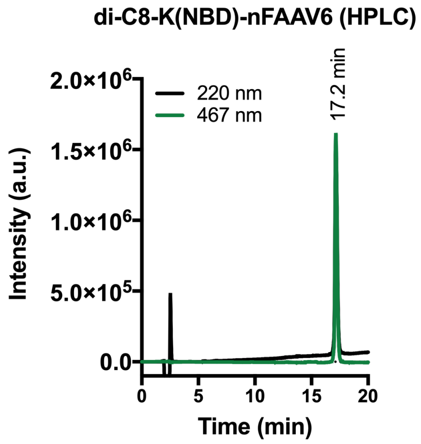


**B**

Calculated values

$[M+H]^+$ = 6086.38

$[M+4H]^{4+}$ = 1522.35

$[M+5H]^{5+}$ = 1218.08

$[M+6H]^{6+}$ = 1015.23

$[M+7H]^{7+}$ = 870.34

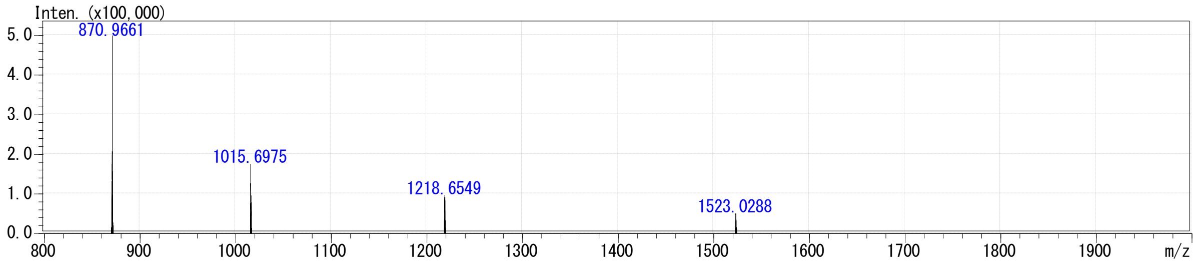


**Figure S7. HPLC profile and MS analysis of 7 [di-C8-K(NBD)-nFAAV6].**

The HPLC profile (A) and the ESI-MS analytical data (B) of the purified peptide. (A) The purity and the retention time were analyzed by RP-HPLC an InterSustainSwift 5$C_{18}$-AR-II column (4.6 mm I.D. × 150 mm) using a linear gradient from 30 to 80% acetonitrile in 0.1% aqueous TFA for 20 min at 40 ºC at a flow rate of 1.0 mL/min. The purity of the peptide was calculated to be over 95% based on the peak area. (B) The calculated masses of multivalent ions are described on top of the observed *m/z* data.

**8** (di-biotin-nFAAV5)

**biotin-DKBLLKXLNKBTDBLSKX-GSGS-C-NH$_2$**
**|**
**biotin-DKBLLKXLNKBTDBLSKX-GSGS-C-NH$_2$**

**A**

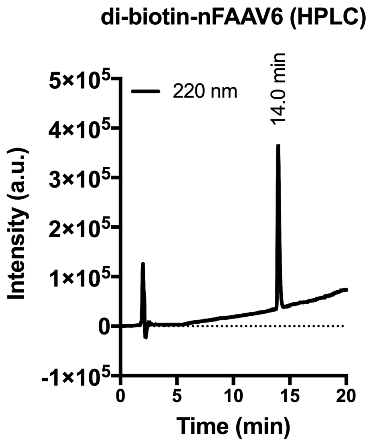


**B**

Calculated values

$[M+H]^+$ = 5223.87

$[M+4H]^{4+}$ = 1741.96

$[M+5H]^{5+}$ = 1306.72

$[M+6H]^{6+}$ = 1045.58

$[M+7H]^{7+}$ = 871.49

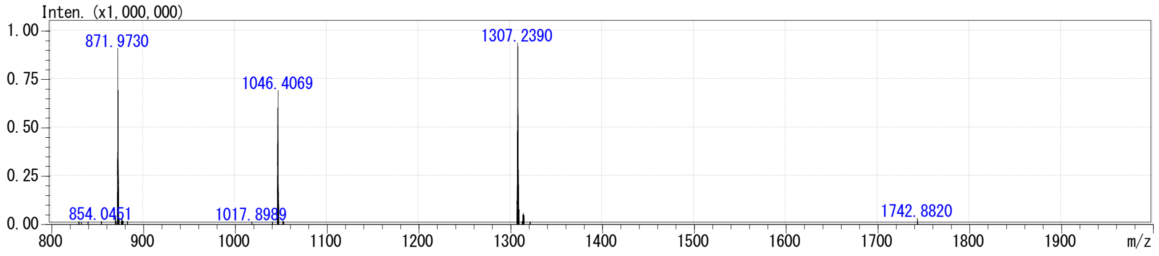


**Figure S8. HPLC profile and MS analysis of 8 (di-biotin-nFAAV5).**

The HPLC profile (A) and the ESI-MS analytical data (B) of the purified peptide. (A) The purity and the retention time were analyzed by RP-HPLC on an InterSustainSwift 5C$_{18}$-AR-II column (4.6 mm I.D. × 150 mm) using a linear gradient from 30 to 80% acetonitrile in 0.1% aqueous TFA for 20 min at 40 ºC at a flow rate of 1.0 mL/min. The purity of the peptide was calculated to be over 95% based on the peak area. (B) The calculated masses of multivalent ions are described on top of the observed *m/z* data.

**9** (biotin-SAH-p53-4)

**biotin-LSQETFZ’DLWKLLZEN-NH$_2$**

**A**

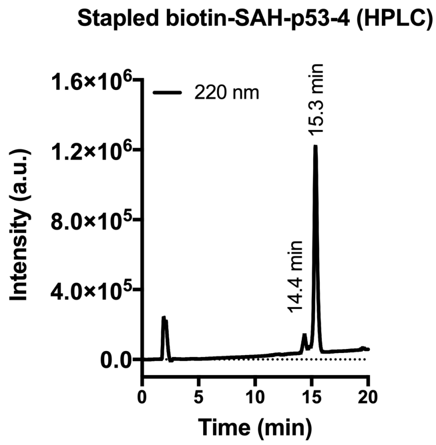


Calculated values

$[M+H]^+$ = 2253.20

$[M+2H]^{2+}$ = 1127.10

$[M+3H]^{3+}$ = 751.74

**B**

Peak at 14.4 min

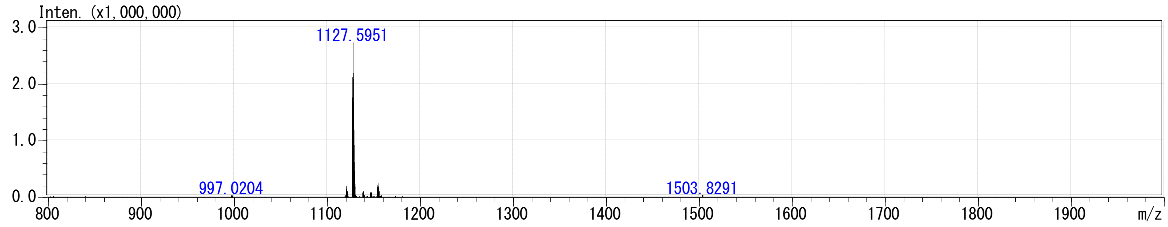


Peak at 15.3 min

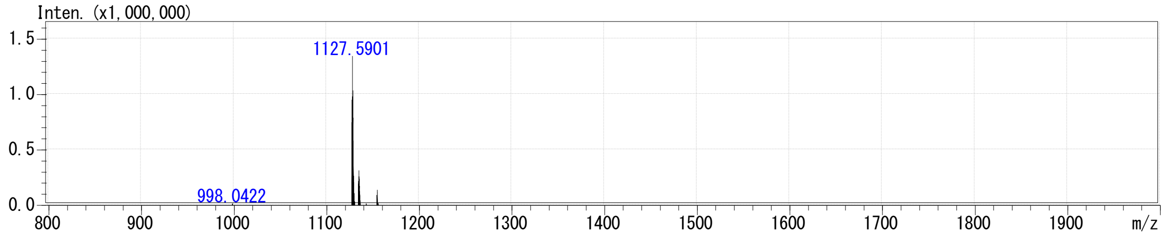


**Figure S9. HPLC profile and MS analysis of 9 (biotin-SAH-p53-4).**

The HPLC profile (A) and the ESI-MS analytical data (B) of the purified peptide. (A) The purity and the retention time were analyzed by RP-HPLC on an InterSustainSwift 5C$_{18}$-AR-II column (4.6 mm I.D. × 150 mm) using a linear gradient from 30 to 80% acetonitrile in 0.1% aqueous TFA for 20 min at 40 °C at a flow rate of 1.0 mL/min. The purity of the peptides in mixture was calculated to be over 95% based on the peak area. (B) The calculated masses of multivalent ions are described on top of the observed *m/z* data.

**10** (biotin-BIM BH3)

**biotin-IWIAQELRRIGDEFNZYYAZR-NH$_2$**

**A**

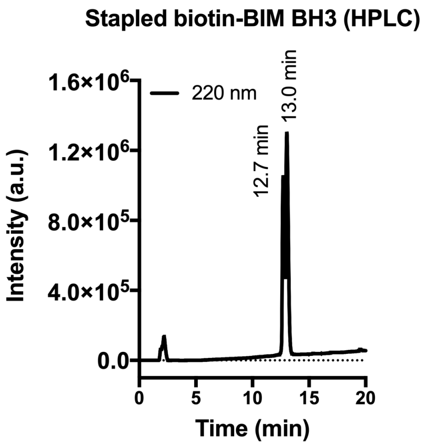


Calculated values

$[M+H]^+$ = 2888.50

$[M+2H]^{2+}$ = 1444.76

$[M+3H]^{3+}$ = 963.51

**B**

Peak at 12.7 min

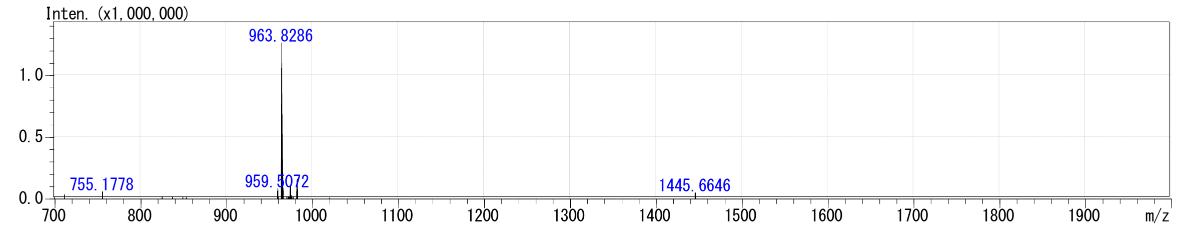


Peak at 13.0 min

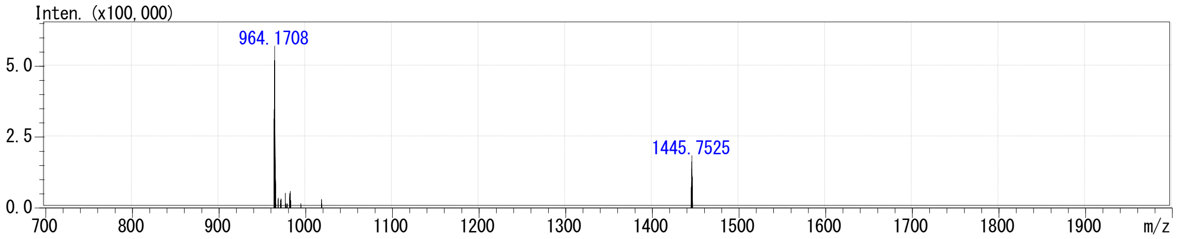


**Figure S10. HPLC profile and MS analysis of 10 (biotin-BIM BH3).**

The HPLC profile (A) and the ESI-MS analytical data (B) of the purified peptide. (A) The purity and the retention time were analyzed by RP-HPLC on an InterSustainSwift 5C$_{18}$-AR-II column (4.6 mm I.D. × 150 mm) using a linear gradient from 30 to 80% acetonitrile in 0.1% aqueous TFA for 20 min at 40 ºC at a flow rate of 1.0 mL/min. The purity of the peptides in mixture was calculated to be over 95% based on the peak area. (B) The calculated masses of multivalent ions are described on top of the observed *m/z* data.

#### Figure S11

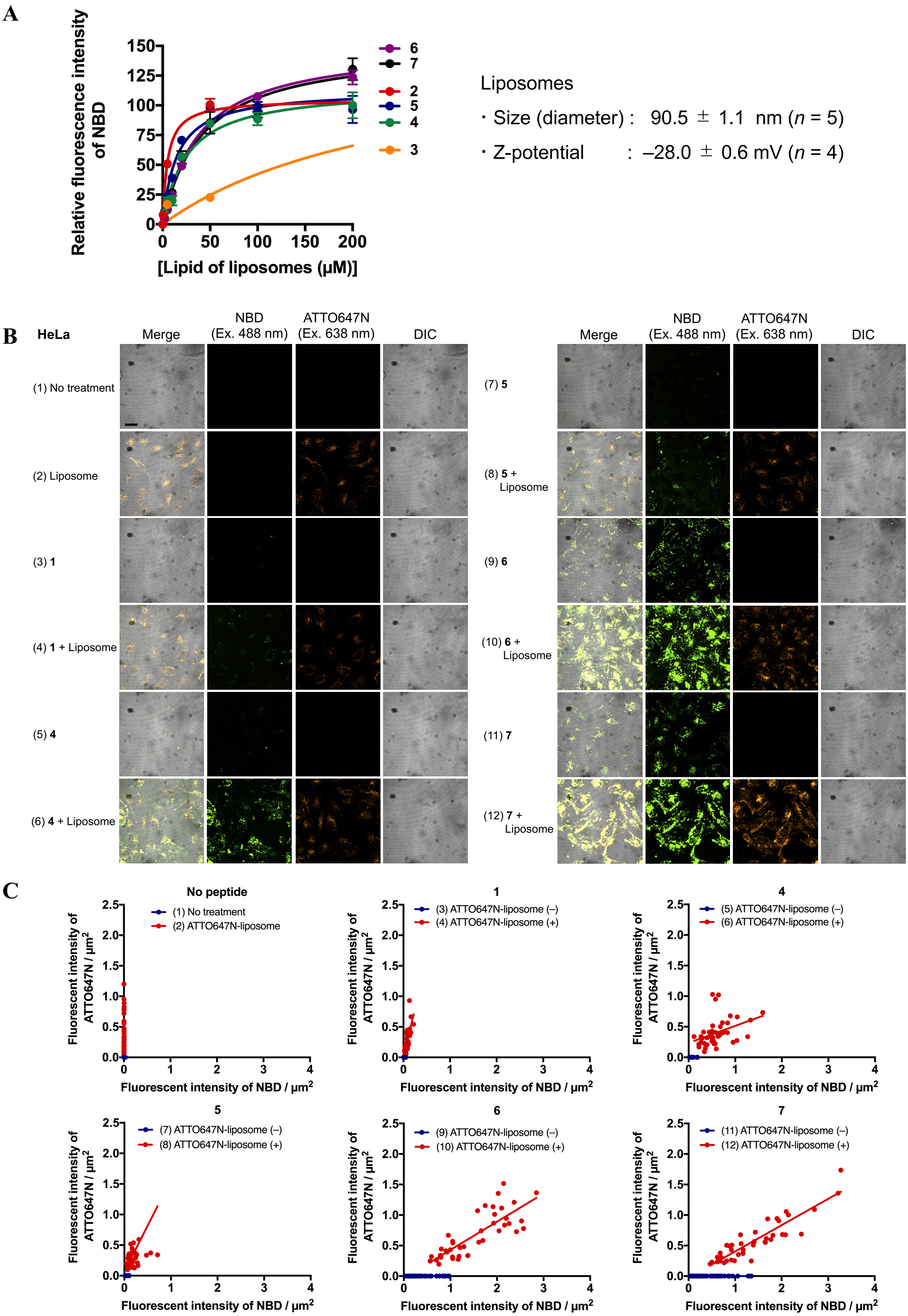
A
Relative fluorescence intensity of NBD
150
125
100
75
50
25
0
0
50
100
150
200
[Lipid of liposomes (μM)]
6
7
2
5
4
3
Liposomes
・Size (diameter) : 90.5 ± 1.1 nm (n = 5)
・Z-potential : –28.0 ± 0.6 mV (n = 4)
B
HeLa
Merge
NBD (Ex. 488 nm)
ATTO647N (Ex. 638 nm)
DIC
(1) No treatment
(2) Liposome
(3) 1
(4) 1 + Liposome
(5) 4
(6) 4 + Liposome
(7) 5
(8) 5 + Liposome
(9) 6
(10) 6 + Liposome
(11) 7
(12) 7 + Liposome
C
No peptide
(1) No treatment
(2) ATTO647N-liposome
1
(3) ATTO647N-liposome (–)
(4) ATTO647N-liposome (+)
4
(5) ATTO647N-liposome (–)
(6) ATTO647N-liposome (+)
5
(7) ATTO647N-liposome (–)
(8) ATTO647N-liposome (+)
6
(9) ATTO647N-liposome (–)
(10) ATTO647N-liposome (+)
7
(11) ATTO647N-liposome (–)
(12) ATTO647N-liposome (+)
Fluorescent intensity of ATTO647N / μm²
Fluorescent intensity of NBD / μm²
0.0
0.5
1.0
1.5
2.0
2.5
0
1
2
3
4

**D**

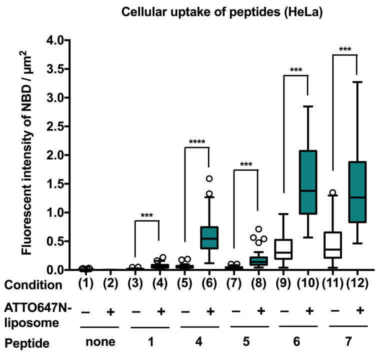


**E**

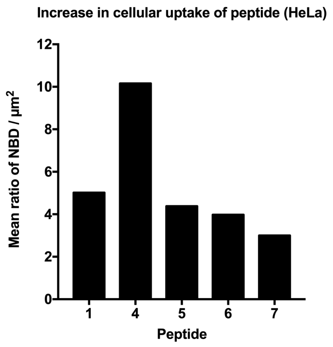


**F**

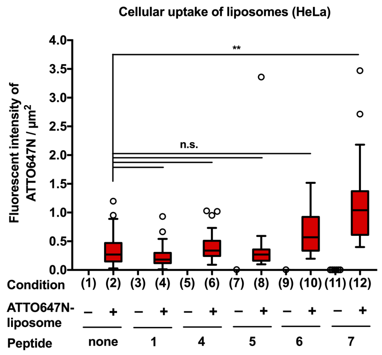


**Figure S11. Cellular uptake of NBD-labeled peptides using ATTO647N-liposomes (HeLa).** (A) The binding saturation curves to sEV-mimicking liposomes for the NBD-labeled peptides, **2**–**7** (mean ± S.D., $n$ = 3). Non-labeling liposomes (POPC:Chol:POPS = 75:15:10 mol%) were used here at the lipid concentrations of 0, 2, 5, 10, 20, 50, 100, 200 µM. The size and Z-potential determined by DLS are shown at the right side. The peptide concentration was fixed at 0.5 µM. (B) The CLSM images of HeLa cells under the conditions of (1)–(12) as shown in the panel. [ATTO647N-liposomes (POPC:Chol:POPS:ATTO647N-DPPE = 75:15:10:0.1 mol%)] = 20 µM at a lipid concentration; [**1**] = 1.2 µM (condition 3) and 2.0 µM (condition 4); [**4**–**7**] = 0.5 µM (conditions 5, 7, 9, and 11) and 1.0 µM (conditions 6, 8, 10, and 12). The concentrations of the peptides alone (conditions 3, 5, 7, 9, and 11) reflect those of the free peptides in the presence of the liposomes (conditions 4, 6, 8, 10, and 12), which were estimated from the liposome binding curves (*refer the Materials and Methods*). The scale bar is 10 µm. (C) Correlation analysis of cellular uptake of the peptide and liposomes. The fluorescent intensities of ATTO647N per cell area (µm$^2$) are plotted as a function of those of NBD per cell area (µm$^2$). (D) Cellular uptake of the peptides in the absence and presence of the liposomes. (E) An increase rate in cellular uptake of the peptides in the presence of the liposomes. The ratios were obtained by dividing the mean values of the fluorescent intensity of NBD / µm$^2$ in the presence of liposomes (even number conditions) by those in the absence of the liposomes (odd number conditions) in (D). (F) Cellular uptake of the liposomes in the absence and presence of the peptides. In the panels (D) and (F), the statistical analysis was performed using $t$-Test (paired two sample for means) with an alpha level of 0.05 ($n$ = 40 cells, *$p < 0.05$, **$p < 0.01$, ***$p < 0.001$, ****$p < 0.0001$).

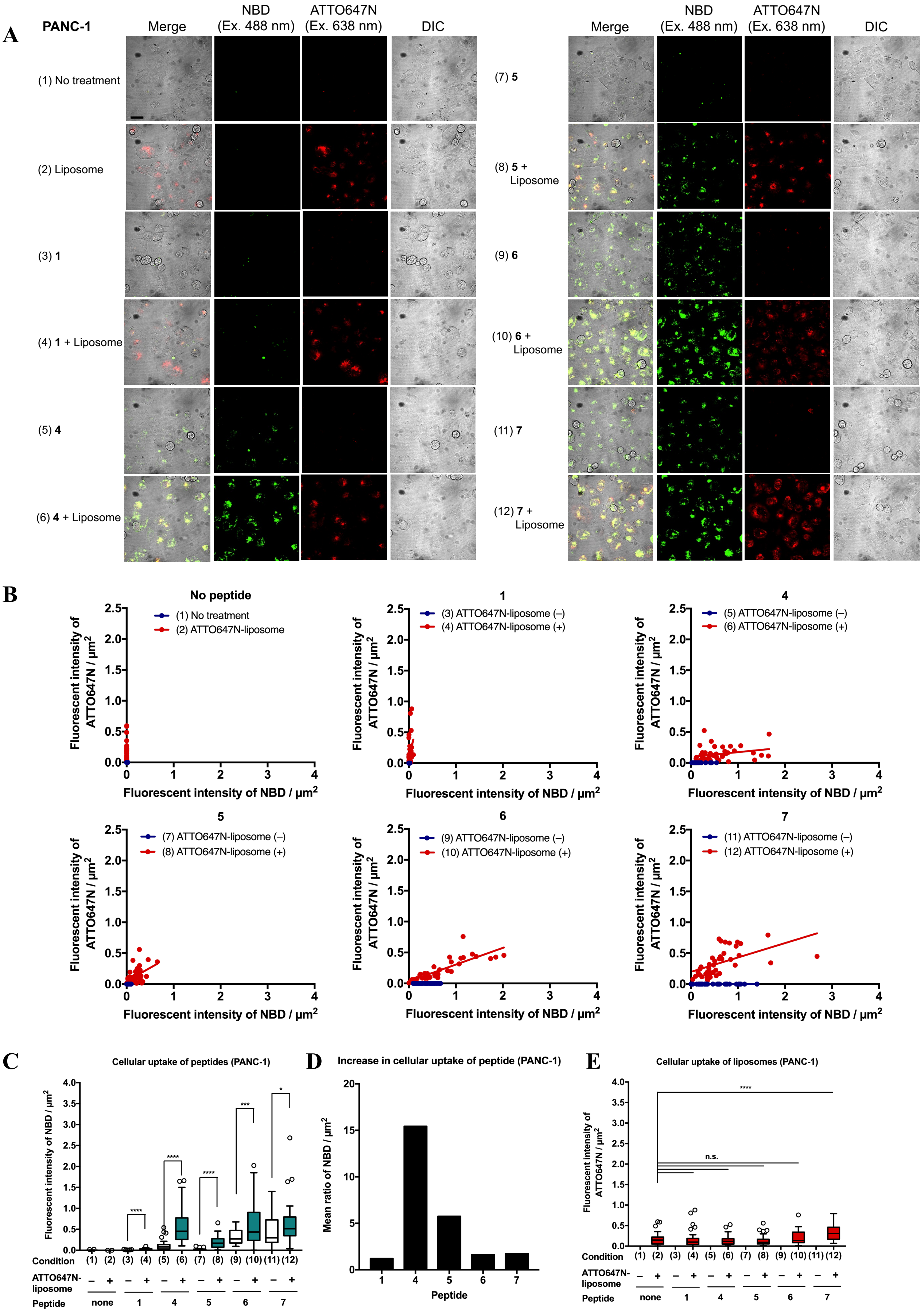


**Figure S12. Cellular uptake of NBD-labeled peptides using ATTO647N-liposomes (PANC-1).** Refer to the legend in Figure S11.

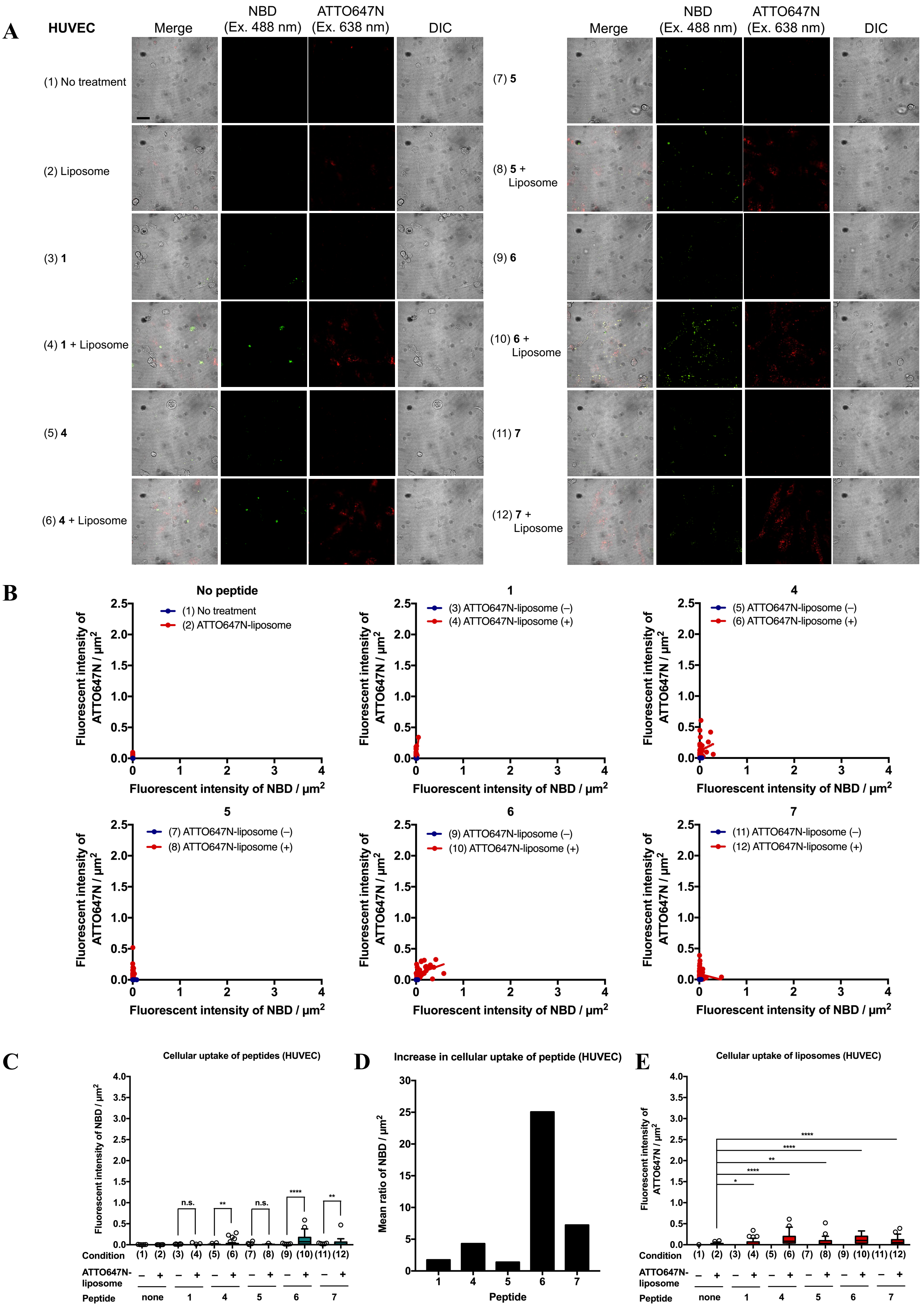


**Figure S13. Cellular uptake of NBD-labeled peptides using ATTO647N-liposomes (HUVEC).** Refer to the legend in Figure S11.

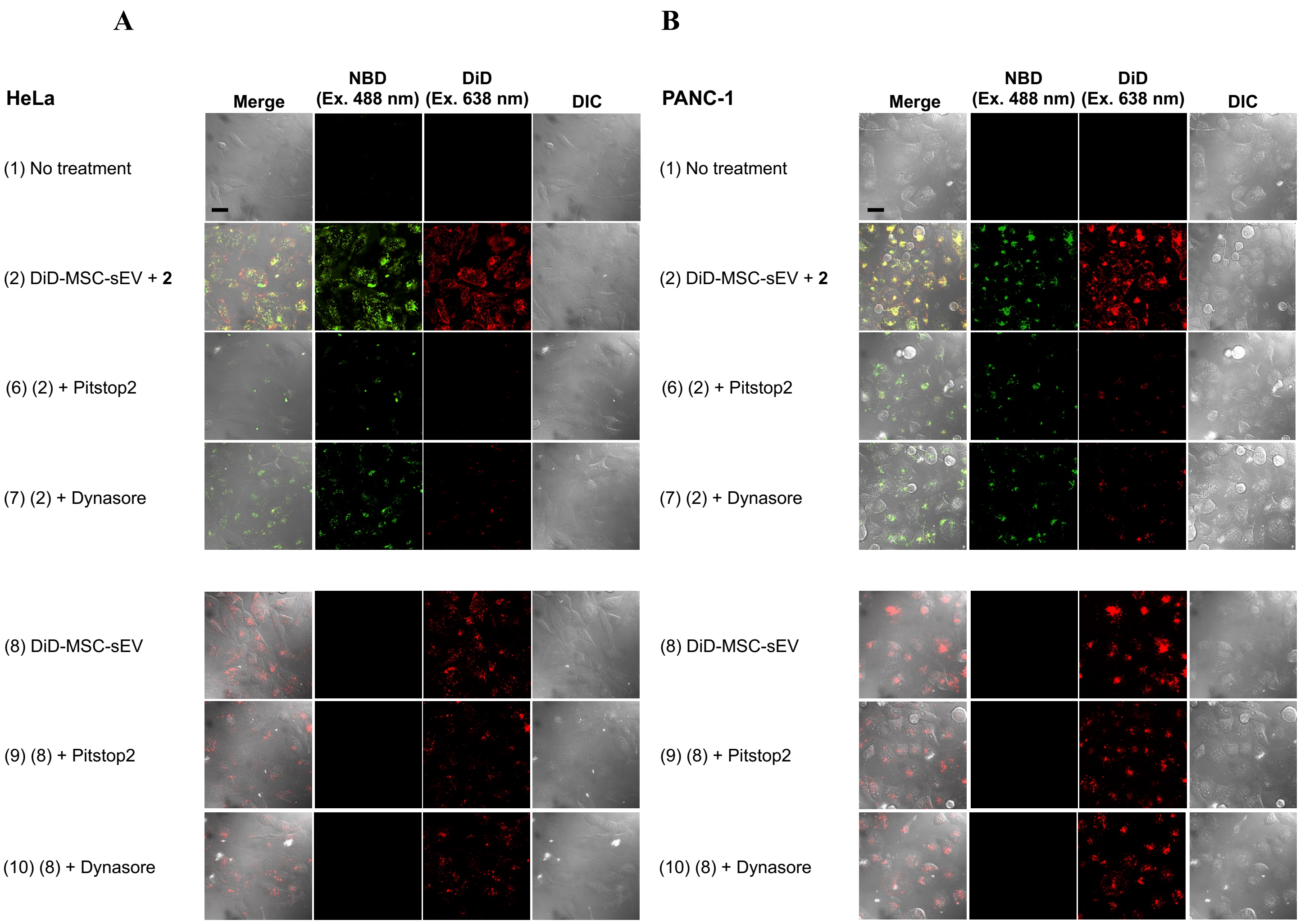


**Figure S14. Cellular uptake of DiD-MSC-sEVs in the presence of inhibitors.**

The representative CLSM images of HeLa (A) and PANC-1 cells (B) under the conditions as shown in the panels. The concentration of the peptide **2** was used at the fincal concentrations of 1 μM. DiD-MSC-sEVs were added to the cells at the fincal concentrations of 1 × $10^8$ and $10^9$ particles/mL in the presence and absence of the peptide **2**, respectively. Because sEVs alone were poorly taken up by cells, a high concentration of sEVs was required for the observation. The scale bar is 10 μm.

A

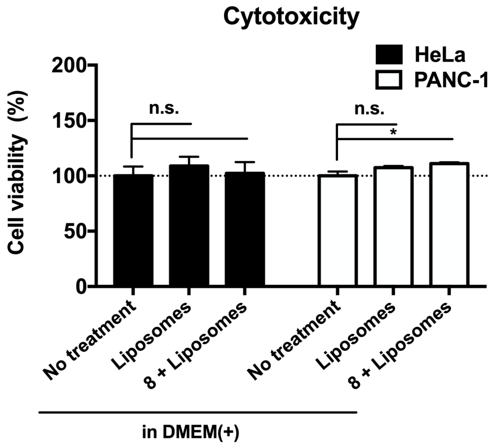


B

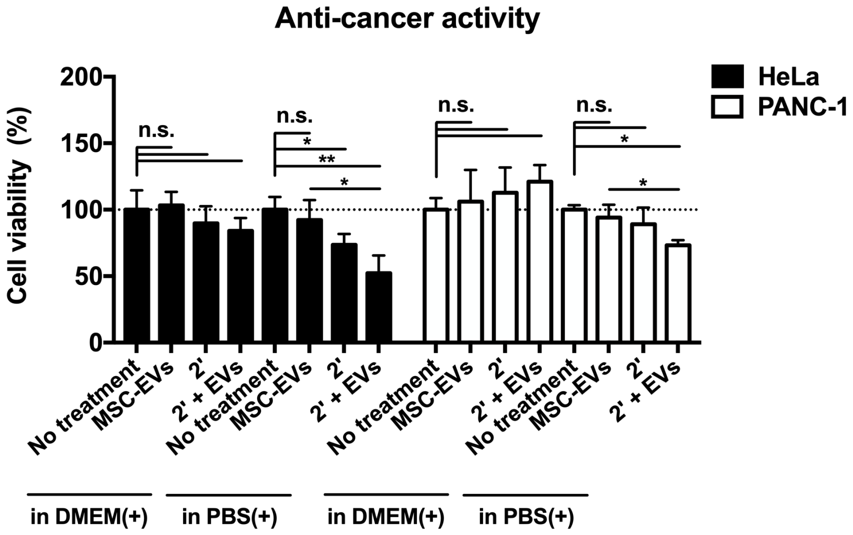


**Figure S15. Cytotoxicity of peptides and anti-cancer activity of MSC-sEVs.**

(A) The cytotoxiticy of the peptide **8** against HeLa and PANC-1 cells in the presence of non-labeling liposomes (30 μM). Because the peptide **8** alone has little effect on cell membranes, its cytotoxiticy can be assessed only in the presence of the liposomes. (B) Anti-cancer activity of non-labeling MSC-sEVs (1 × $10^9$ particles / mL) against HeLa and PANC-1 cells in the presence of the peptide **2'** (1 μM) in DMEM(+) or PBS(+). In the panels (A) and (B), the statistical analysis was performed using *t*-Test (paired two sample for means) with an alpha level of 0.05 ($n = 3$, $*p < 0.05$, $**p < 0.01$, $***p < 0.001$, $****p < 0.0001$).

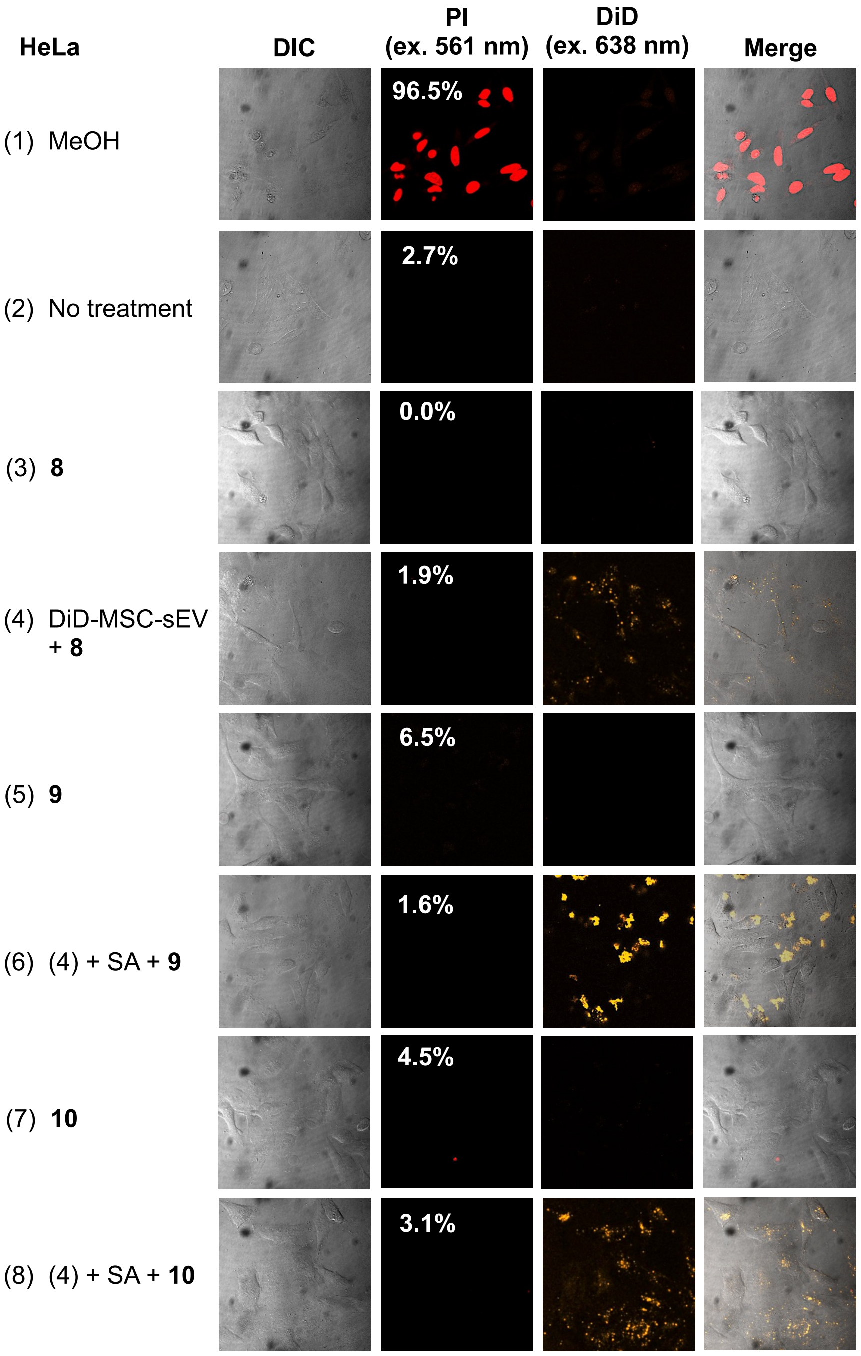


**Figure S16. Membrane perturbation assay (HeLa).**

Membrane perturbation assay of the drug delivery system using a membrane-impermeable dye, PI. The percentage indicates the ratio of the dead cells, which were stained by PI ($n$ = 37–65 cells). The cells treated with MeOH was used as a control.

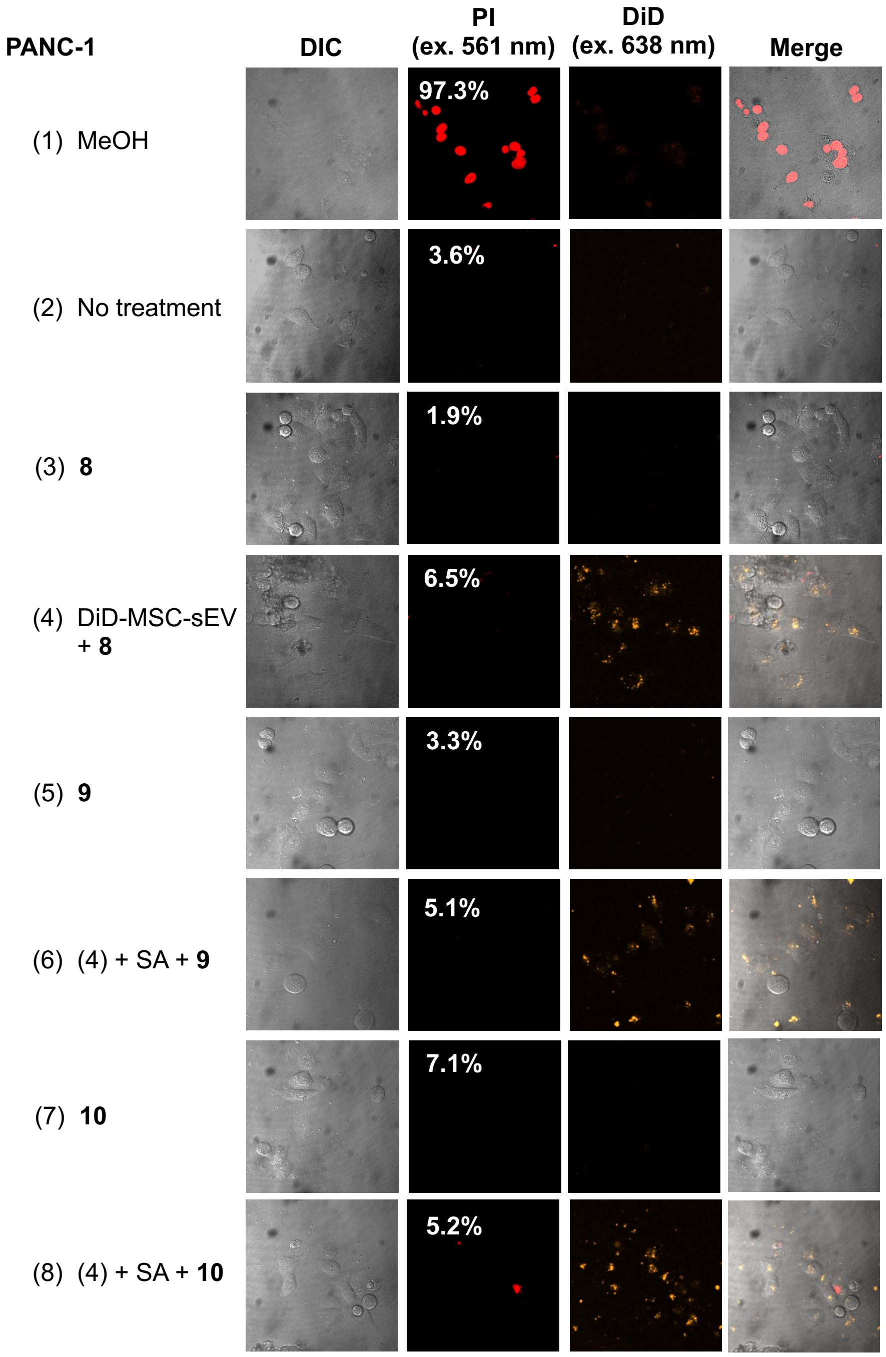


**Figure S17. Membrane perturbation assay (PANC-1).**

Membrane perturbation assay of the drug delivery system using a membrane-impermeable dye, PI. The percentage indicates the ratio of the dead cells, which were stained by PI ($n$ = 28–58 cells). The cells treated with MeOH was used as a control.
